# Change in topological linking number during Xer recombination at the plasmid pSC101 *psi* site

**DOI:** 10.64898/2026.02.23.706985

**Authors:** James I. Provan, Anna O. Tolmatcheva, David J. Sherratt, Sean D. Colloms

## Abstract

The XerCD recombinase functions with accessory proteins PepA, ArcA and ArgR at plasmid recombination sites such as *psi* and *cer* to ensure stable monomeric plasmid inheritance. Xer recombination acts only on directly repeated *psi* sites, and recombination produces a specific catenane of two product circles interlinked exactly four times. Here we measure the precise change in topological linkage (ΔLk) that occurs during Xer recombination at *psi*. We use a DNA substrate with close-spaced *psi* sites that recombines to produce one circle of 398 bp and another of 3039 bp and demonstrate that the small circle is exclusively the -1 topoisomer. Using a purified topoisomer of the substrate, we show that Xer recombination proceeds with a linkage change (ΔLk) of +4. Similar experiments using a substrate with equally spaced *psi* sites agreed with this result. The measured linkage change is consistent with a reaction mechanism for tyrosine recombinases in which the sites align antiparallel prior to recombination and recombine *via* a Holliday junction intermediate. Four negative supercoils are converted to catenation nodes by strand exchange, providing an energetic driving force for the reaction. We compare this result to the mechanisms of serine recombinases and the recently discovered bridge RNA-guided recombinases.

**Graphical Abstract:** 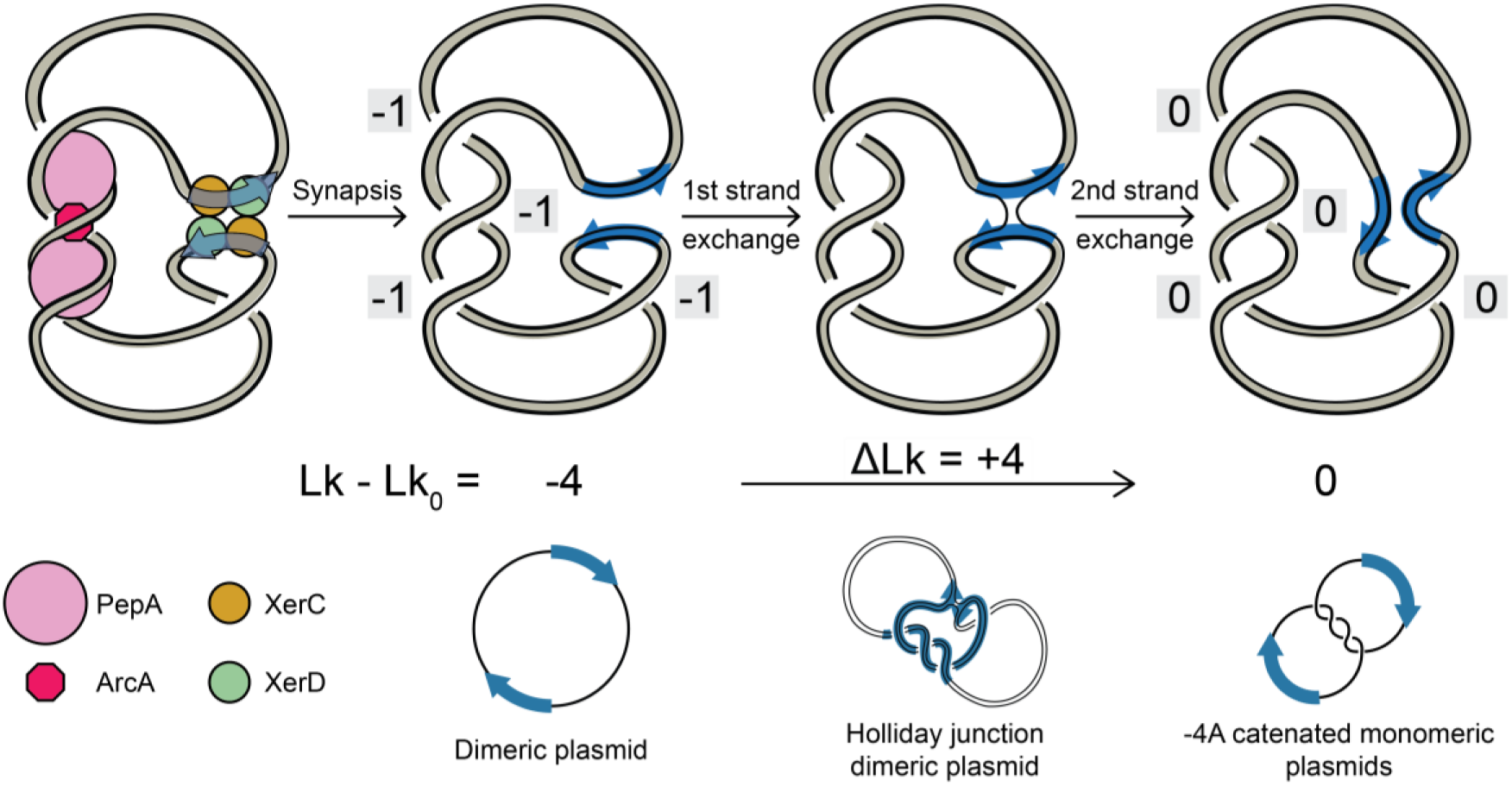

## INTRODUCTION

Site-specific recombinases mediate programmed DNA rearrangements by binding to short DNA recombination “sites”, cutting the DNA strands, and then re-joining them in a recombinant configuration. The biological roles of site-specific recombination include insertion and excision of bacteriophage and other mobile elements from the genome of their host ^1–6^, regulation of gene expression by DNA inversion ^7–10^, hoarding potentially useful gene collections for later use in structures known as integrons ^11^, resolution of transposition intermediates ^12^, and DNA multimer resolution systems of viral elements, plasmids, and circular chromosomes ^13–17^.

Site-specific recombination is generally initiated by the recombinase protein binding to imperfect dyad symmetric sites as dimers and then tetramerizing to bring two sites together in a synaptic complex. This is followed by cleavage and re-joining of all four DNA strands near the centres of the sites to form recombinant products in which the left half of each site is joined to the right half of the other. There are two major families of site-specific recombinase: serine recombinases use a serine hydroxyl group to cleave the DNA by nucleophilic attack on the DNA backbone, while tyrosine recombinases utilise a tyrosine hydroxyl group for the same purpose. Serine recombinases cut all four strands of the participating DNA sites before rotating one pair of half sites by 180° relative to the other and re-joining all four strands (Figure 1a-i) ^18–21^. Tyrosine recombinases cleave and re-join one pair of strands to form a Holliday junction, then cleave and re-join the other pair to resolve the Holliday junction forming the recombinant product (Figure 1a-ii) ^22–26^. Well known tyrosine recombinases include Cre, FLP, λ integrase, and the Xer recombination system. Recently the PIV invertases of *Moraxella* species and the IS110 family transposases have been found to make up a third family of site-specific recombinases. These enzymes catalyse site-specific recombination via a Holliday junction intermediate using a serine nucleophile, four acidic residues (DDED) to coordinate a divalent metal ion, and a bridge RNA involved in site recognition ^27^.

**Figure 1:**
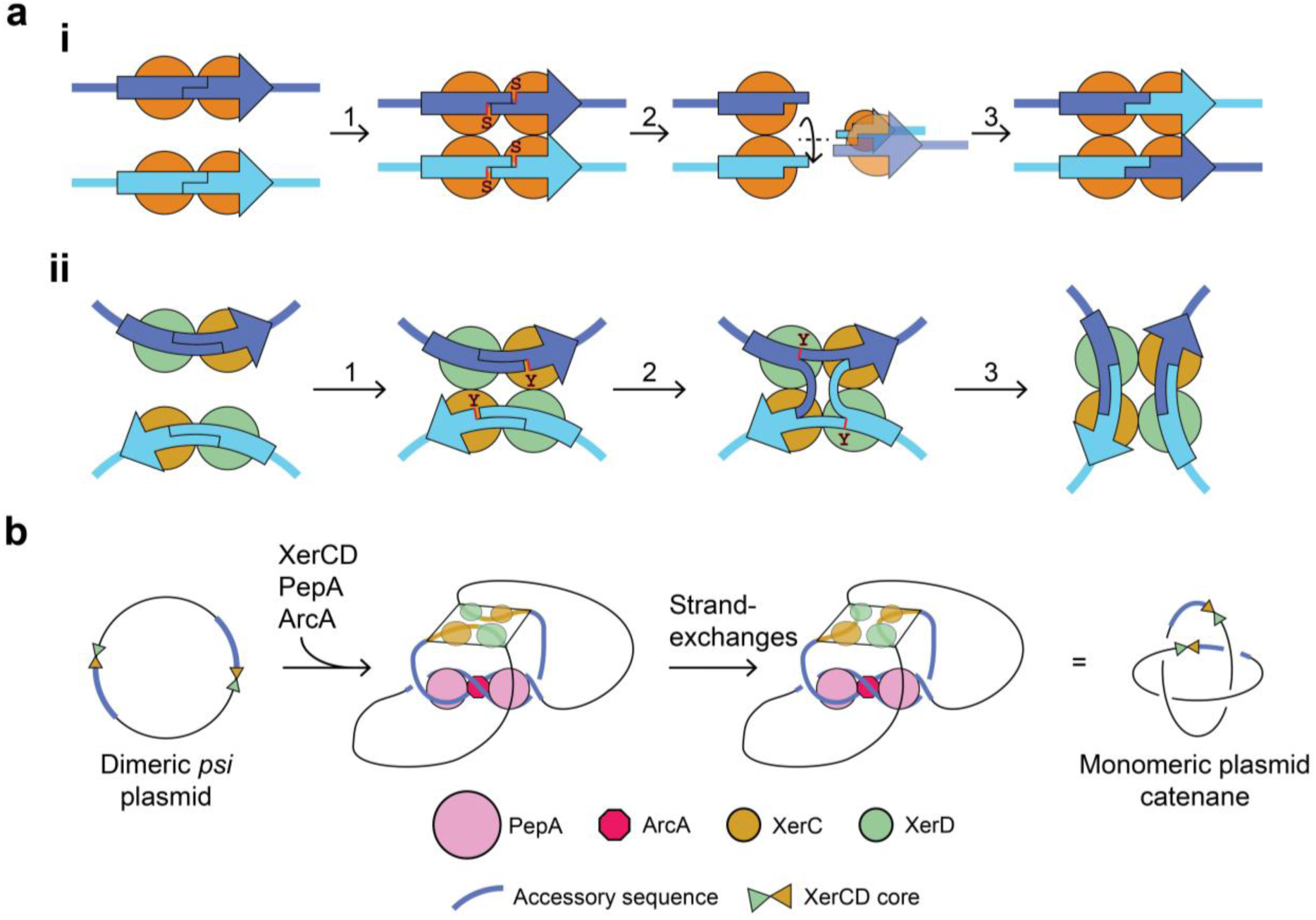
Mechanisms of site-specific DNA recombination and the topology of Xer recombination at *psi*. **a, i**, Serine recombinases bind their recombination sites as dimers and tetramerise with sites aligned in parallel (1). Active-site serine residues cleave all four strands (2), followed by a 180° subunit rotation of one side of the tetramer compared to the other (3). The four strands are re-ligated, forming two recombinant sites each composed of two different original half-sites. **ii,** Tyrosine recombinases bind to their recombination sites as dimers (1) and tetramerise with sites aligned in an approximately anti-parallel orientation. Active-site tyrosine residues on one pair of recombinases catalyse pairwise strand-cleavage and transfer (2) forming a Holliday-junction intermediate. Isomerisation of the HJ tetramer places the other pair of recombinases in the activated conformation for cleavage and strand exchange of the other pair of strands (3), resulting in two recombinant sites. **b,** The topological model of plasmid dimer resolution by Xer recombination at pSC101 *psi.* On a plasmid dimer, *psi* accessory sequences wrap around the accessory proteins PepA and ArcA, trapping three right-handed crossings and aligning the XerCD core recombination sites for XerC-first activity. Strand exchange *via* a Holliday junction produces the observed right-handed four-node catenane, that can be separated into component rings by a type II topoisomerase.

Xer site-specific recombination systems are found in all bacterial species with circular chromosomes and function to ensure faithful segregation of the chromosome at cell division ^13^. Double-strand break repair by homologous recombination at broken replication forks and other DNA damage generates crossovers. Following the replication of circular chromosomes, an odd number of crossovers will fuse the two chromosomes into a double sized dimeric circle that cannot be segregated to both daughter cells at cell division without DNA breakage ^17,28^. Xer recombination, at a site termed *dif* in the replication terminus region of the bacterial chromosome, resolves these chromosome dimers into two separate chromosomal monomers which can be segregated accurately to both daughter cells ^29,30^. The *dif* site consists of 11 bp binding sites for XerC and XerD separated by a 6 bp spacer region. The XerC and XerD tyrosine recombinases bind to two *dif* sites to form a heterotetrameric synaptic complex, then cleave on either side of the 6 bp spacer to catalyse recombination *via* a Holliday junction intermediate (Figure 1aii) ^30–34^. Xer recombination at *dif* is activated by the DNA translocase FtsK only after FtsK has pumped the two monomeric halves of the chromosomal dimer into the nascent daughter cells, thus ensuring accurate chromosome segregation ^35–40^.

Multicopy plasmids in bacteria are also subject to destabilising multimer formation ^41,42^. Inter-plasmid homologous recombination and/or rolling circle replication produces plasmid dimers and higher multimers; that is, circular concatemers consisting of 2 or more copies of the plasmid sequence joined in direct (head to tail) repeat ^41,43^. Plasmid multimers are generally maintained at a lower copy number than monomers because replication initiation is negatively regulated *via* origin-encoded repressive mechanisms, which ultimately regulate the number of origins present without sensitivity to the number of independently replicating circular copies^44^. Therefore, plasmid dimers are maintained at approximately half the copy number of monomers, tetramers at a quarter the copy number, etc. This reduction in the number of independently segregating plasmid copies causes an increase in the chance of producing plasmid-free cells at cell division, leading to plasmid segregational instability. To counter this, many multicopy plasmids contain recombination sites that utilise the chromosomally encoded Xer recombinases to convert plasmid multimers back to monomers ^45–49^.

Plasmid sites for Xer recombination, such as *cer* from plasmid ColE1 and *psi* from pSC101, contain a *dif*-like core adjacent to accessory DNA sequences that are bound by accessory proteins to regulate the directionality of recombination. The architectural DNA binding protein PepA is the main accessory protein for Xer recombination at *cer* and *psi* ^50–53^. Along with the secondary accessory protein (the transcriptional regulators ArgR for *cer*-like sites or ArcA for *psi*-like sites), PepA wraps the accessory sequences of two recombination sites plectonemically around each other approximately 3 times (Figure 1b) ^47,54–56^. Formation of this interwrapped complex activates strand exchange by the Xer recombinases at the *dif*-like core sites, and is favoured between directly repeated sites in a negatively supercoiled DNA molecule but not between sites in inverted repeat or on separate DNA molecules ^51^. This ensures that recombination at *cer* and *psi* occurs only between direct repeat sites on the same supercoiled DNA molecule ^14,47^ so that multimers are resolved to monomers, but intermolecular recombination to form multimers does not occur. Recombination takes place in a specific synapse with the accessory sequences interwrapped a fixed number of times and strand exchange occurs by a defined mechanism so the product has a specific topology - a 4-node right-handed catenane^57^ with antiparallel *psi* sites (Figure 1b) ^14^.

The exact mechanism of synapsis and strand exchange by tyrosine recombinases has been the subject of debate. Early models for recombination by tyrosine recombinases proposed that sites were aligned in parallel throughout the reaction ^58,59^. Electron microscopy of complexes formed between the FLP recombinase and its recombination site suggested a strong preference for parallel site alignment ^60^. Structural studies on Cre, lambda integrase and Xer recombinases all support the presence of an antiparallel square-planar Holliday junction reaction intermediate ^24,25,61,62^, while topological studies and ring closure experiments support the presence of a chiral right-handed initial synapse ^63,64^. Measuring the change in DNA linking number during the reaction may help to distinguish between these mechanistic differences. Such studies have been performed for Tn3 resolvase, a member of the serine recombinase family ^19,65^. However, recombination by most members of the tyrosine recombinase family (e.g. Cre and lambda integrase) catalyse strand exchange after random collision of their bound recombination sites, which traps random numbers of supercoils between the recombining sites and yields knots and catenanes of variable topologies. For this reason, it has been difficult to determine a single DNA linkage change associated with tyrosine recombinase activity and to pin down the exact topology of the reaction. Here we use the fixed topology of Xer recombination at *psi* to show that the reaction takes place with a specific linkage change fully consistent with sequential strand exchange of sites in an antiparallel arrangement, *via* a Holliday junction intermediate. The conversion of four negative supercoils to more favourable catenation crossings during the reaction drives the reaction forward, making it essentially unidirectional.

## MATERIALS AND METHODS

### Plasmids and strains

pSDC133 (3046 bp) contains a single pSC101 *psi* site in the pUC18 multicloning site (MCS), while pLN5 (3046 bp) is pSDC133 mutagenized to contain a MluI restriction site within *psi*. pCLOSE (pSDC153, 3437 bp) is pSDC133 with a second *psi* site inserted within the MCS in the same orientation ^47^. pCLOSEΔ (3039 bp) is the product of *in vivo* Xer recombination of pCLOSE in the Xer^+^ *E. coli* strain DS941. πAN7 is an 885 bp microplasmid that uses a *sup^F^* amber-suppressor tRNA as selection marker that requires the amber-codon mutated ampicillin/tetracycline resistance plasmid “p3” (60 kb) ^66,67^. pEVENΔ (1260 bp) contains the EcoRI-SalI *psi* fragment of pSDC133 subcloned into the MCS of πAN7, while pEVENΔ*-*MluI is the same construction derived from pLN5. An equally spaced *psi* recombination substrate, pEVEN, was constructed by linearising both pEVENΔ and pEVENΔ*-*MluI with EcoRI, ligating the resultant fragments together using T4 DNA ligase, and selecting the correct pseudo dimeric plasmid containing one *psi* and one *psi*-MluI site. Plasmid constructions were verified before use by analytical restriction digests and Sanger sequencing. Plasmid sequences are available in the Supplementary Materials.

All strains used in this work were derivatives of *E. coli* K-12 strain DS941 ^68^ (AB1157 *recF lacI^q^*, *lacZΔM15*). pUC18-derived plasmids containing *psi* were maintained in DS984 (DS941 *xerC::CmR*), while pEVENΔ and derivatives were maintained in a *sup*^0^ *xerC^-^* derivative of DS941 (DS953 *xerC::CmR*) containing the p3 selection plasmid.

### Commercial Enzymes

Restriction enzymes, nicking endonucleases, RecJ_f_ and λ-exonuclease were obtained from New England Biolabs and used in Cutsmart buffer. Calf thymus topoisomerase I (TOP1) was obtained from Invitrogen (ThermoFisher) and *E. coli* topoisomerase IV (TOP4) was obtained from Inspiralis. Topoisomerase I reactions were carried out in 50 mM Tris-HCl pH 7.5, 50 mM KCl, 10 mM MgCl_2_, 0.5 mM dithiothreitol, 0.1 mM EDTA and 30 μg/ml bovine serum albumin (TOP1 buffer), while topoisomerase IV reactions were in 50 mM Tris-HCl pH 7.5, 100 mM potassium glutamate, 6 mM MgCl_2_, 10 mM dithiothreitol, 1.0 mM ATP and 50 μg/ml bovine serum albumin (TOP4 buffer).

### Protein purification

Wild-type XerC (Uniprot: P0A8P6), an MBP-XerC fusion protein, wild-type XerD (Uniprot: P0A8P8), and wild-type PepA (Uniprot: P68767) were overexpressed in *E. coli* and purified to homogeneity as described in references ^31, 33, 69^, and ^57^ respectively. The wild-type XerC protein was used during the 2D agarose gel Xer recombination experiments, whereas MBP-XerC was used during the 1D PAGE Xer recombination experiment. The MBP-XerC fusion protein was used without cleavage of the fused MPB solubility tag. For each protein purification, the peak absorbance fractions of their final FPLC step were pooled and dialysed against a storage buffer containing 50 mM Tris-HCl pH 8.0, 1 M NaCl, 1 mM EDTA, 2 mM TCEP, and 50% glycerol. Proteins were routinely stored at -20°C and prediluted prior to use with a dilution buffer containing 10 mM Tris-HCl pH 7.5, 10 mM MgCl_2_, 75 mM NaCl, 2 mM TCEP, and 50% Glycerol.

### DNA purification

Large scale preparations of supercoiled plasmids were generated by alkaline lysis followed by ethidium bromide / CsCl density gradient centrifugation or using a Maxiprep kit (Qiagen). Where we refer to extraction with phenol:chloroform and precipitation with ethanol, this represents extraction of the sample with an equal volume of phenol:chloroform:isoamyl alcohol (25:24:1), followed by at least two extractions of the aqueous phase with equal volumes of chloroform:isoamyl alcohol (24:1), followed by precipitation of DNA with two volumes of ethanol in the presence of 0.3 M sodium acetate as per standard methods ^70^. DNA was resuspended in TE buffer (10 mM Tris-HCl pH 8.0, 1 mM EDTA).

### *In vitro* Xer recombination reactions

Xer recombination reactions contained a concentration of 21 nM supercoiled substrate plasmid DNA in a buffer containing 50 mM Tris-HCl pH 8.0, 25 mM KCl, 1.25 mM EDTA, 5 mM spermidine, 25 μg/ml BSA, and 10% glycerol. PepA was added to a final concentration of ∼300 nM (as hexamer) and XerC and XerD were added to 250 nM (monomer). Reactions were incubated at 37°C for 1-2 hours and the DNA was purified by extraction with phenol:chloroform and precipitation with ethanol.

### Preparation of topoisomer ladders for 2-dimensional gels

Topoisomers relaxed at different concentrations of intercalator were mixed at suitable ratios to produce marker ladders for 2-dimensional gel electrophoresis. Individual reactions containing 1 μg of pCLOSE, pCLOSEΔ, pEVEN or pEVENΔ were relaxed with calf-thymus topoisomerase I in the presence of 0, 300, 600, or 1000 μg/ml chloroquine or 2, 3 or 4 μg/ml ethidium bromide in 50 μl of TOP1 buffer. Reactions were started by the addition of 2 Units of Topoisomerase I and were incubated at 37°C for 60 minutes. Reactions at different intercalator concentrations were stopped by extraction with phenol, pooled and further extracted with phenol:chloroform followed by ethanol precipitation.

### Purification of single topoisomers for linkage change experiments

Natively supercoiled pCLOSE or pEVEN were loaded in multiple adjacent lanes of 1.2% SeaPlaque low melting point (Lonza) agarose gels in 1 x TAE running buffer (40 mM Tris base, 20 mM acetic acid and 1 mM EDTA) with 1.5 μg/ml chloroquine. After electrophoresis, gels were washed in several changes of 1 x TAE containing 0.03% SDS to remove chloroquine, and stained with ethidium bromide. Individual topoisomers were cut from the gel under long wavelength (360 nm) UV illumination. The DNA was purified by melting the agarose at 65°C followed by immediate addition of an equal volume of phenol. After vigorous mixing and centrifugation, the aqueous phase was removed to a fresh tube and subsequently extracted twice with an equal volume of phenol, then with phenol:chloroform, and finally chloroform. The DNA was recovered by precipitation with ethanol and resuspended in TE.

### Linkage change analysis by 2D gel electrophoresis

Purified topoisomers of pCLOSE or pEVEN were reacted with PepA, XerC, and XerD as described above. After extraction with phenol and precipitation with ethanol, pCLOSE was cut with BamHI to linearise the small product circle, releasing the large pCLOSEΔ-sized circle. Reactions with pEVEN were cleaved with MluI to linearise one circle, releasing the other identically sized circle from the catenane. Substrate and product topoisomers were analysed alongside suitable topoisomer ladders by 2-dimensional gel electrophoresis in large format (∼25 x 25 cm) horizontal 1.2% agarose gels in 1 x TAE. The two dimensions were carried out at two different concentrations of chloroquine as indicated in individual figures. The same concentration of chloroquine was included in the gel and running buffer for each dimension. For gels where the chloroquine concentration was higher in the second dimension than the first, the DNA-containing region of the gel was excised after the first dimension, rotated by 90° and cast into a new agarose gel slab containing the higher chloroquine concentration. For gels in which the second dimension contained no intercalator, the entire gel was washed with gentle agitation in 1x TAE containing 0.03% SDS for 8 hours with three changes of buffer to remove the chloroquine. The intact gel was then put back in the running tank and subjected to electrophoresis at right angles to the original direction in 1 x TAE with 0.03% SDS. Electrophoresis was carried out for 16-20 hours at 50V-80V at room temperature with buffer recirculation. Gels were capillary blotted in neutral conditions onto Hybond-N membrane (Amersham) according to the manufacturer’s instructions. pCLOSE blots were hybridised with ^32^P random-primed EcoRI-digested pSDC133, while pEVEN experiments were probed with ^32^P random-primed MluI-digested pEVENΔ*-*MluI. After high stringency washing, the signal was detected by exposure to X-ray film (Fuji). Blot densitometry of the pEVEN product topoisomers was carried out using the FIJI image analysis package ^71^.

### Determining supercoiling levels of the small circle

Large scale (∼200 μg pCLOSE DNA substrate in a total volume of 5 ml) Xer reactions were performed for subsequent analysis of the small circle product. To produce small circle markers with different supercoiling levels, pCLOSE Xer reactions were treated with Nt.BsmAI to nick the small circle once and the large circle five times. After extraction with phenol:chloroform and recovery of the nicked catenane by ethanol precipitation, the nicks were sealed with T4 DNA ligase at different concentrations of ethidium bromide. Reactions were extracted with phenol:chloroform to remove the ethidium and inactivate ligase, and the DNA was recovered by ethanol precipitation.

The large product circle and unrecombined pCLOSE were specifically depleted from Xer reactions or ligation reactions by one-pot co-incubation with HpyCH4V endonuclease, RecJ_f_ exonuclease, and λ-exonuclease in 1x Cutsmart reaction buffer at 37°C for 3 hours^72^. Depletion reactions contained approximately 1 unit HpyCH4V, 1 unit λ-exonuclease, and 7.5 units RecJ_f_ per μg total DNA. Prior to topoisomerase treatment, nucleases were inactivated by phenol-chloroform extraction and ethanol precipitation. Calf-thymus Topoisomerase I and *E. coli* Topoisomerase IV were used at approximately 1 unit per 100 ng small product circle in the supplied reaction buffers, and incubated at 37°C for 1 hour. Topoisomerases were inactivated through the addition of SDS and Proteinase K (Roche) to 1% and 250 μg/ml final concentrations respectively, followed by incubation at 45°C for 1 hour. To ensure consistent migration in polyacrylamide gels, all samples were adjusted to 1x Cutsmart buffer or 1x TOP1 reaction buffer. NEB Purple Loading Dye was added to 1x concentration prior to loading. Large format (17 x 14 cm x 0.15 cm) vertical gels were prepared containing 5% acrylamide (37.5:1 acrylamide: N,N′-methylenebisacrylamide), 1x TB buffer (89 mM Tris base, 89 mM boric acid), and 10 mM CaCl_2_. 1xTB with 10 mM CaCl_2_ was used as running buffer without recirculation at 95V for 18 hours at room temperature. Gels were stained in 0.5x SYBR Gold (Invitrogen) for 30 minutes, de-stained in deionised water for 10 minutes, and imaged using a Typhoon FLA-9500 laser scanner (473 nm laser, LPB filter, PMT 450V). Densitometry of the small circle products was carried out using the FIJI image analysis package ^71^.

## Results

### Topological terminology and strategy used to determine DNA linkage change

The aim of this work was to measure the change in linking number (ΔLk) of the DNA during Xer recombination in order to provide novel information on the molecular mechanism of the reaction. The linking number (Lk) of a circular DNA molecule is the number of times one strand of the double stranded DNA winds around the other. ΔLk of a site-specific recombination reaction is defined as the difference between the total linking numbers of the product and the substrate DNA molecules, ignoring any catenation or knotting of the products. For a deletion reaction between directly repeated sites where a circular substrate (S) is converted to two circular products (P1 and P2) this gives:

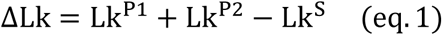

The absolute linking number Lk of a large DNA circle cannot easily be measured, but the linkage difference (Lk – Lk_0_), the number of helical turns relative to the relaxed linking number (Lk_0_) can be measured by gel electrophoresis. On a 2-dimensional gel, with different concentrations of the DNA intercalator chloroquine in both dimensions, individual topoisomers differing from each by just one turn of the helix can be separated. The most relaxed topoisomer in each dimension will be the slowest migrating band in the ladder in that dimension and will run at approximately the same position as the nicked circle of the same size. Using a reference marker ladder containing a range of different topoisomers, the linkage difference can be counted relative to the most relaxed topoisomer (that with Lk ≈ Lk_0_). Note that other authors have previously used the term ΔLk to refer to linkage difference (loosely the amount of supercoiling in the molecule). Herein we refer to the linkage difference as (Lk *–* Lk_0_) throughout, and reserve the term ΔLk for the change of linking number during recombination.

Lk_0_ is related to the relaxed helical repeat of the DNA (H_0_; the average number of bp per turn of DNA; approximately 10.5 bp/turn for standard B-form DNA) as follows:

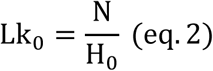

where N is the length of the DNA in bp. Note that while Lk can only take integer values, Lk_0_ is usually non integral and dependent on the conditions of the solute (pH, temperature, etc).

Assuming that both products (P1 and P2) of a site-specific recombination reaction between direct repeat sites have the same relaxed DNA helical repeat as the substrate, then because the length of the substrate is the sum of the lengths of the two products, the relaxed linkage of the two products (Lk_0_^P1^ and LK_0_^P2^) add up to the relaxed linkage of the substrate (Lk_0_^S^).

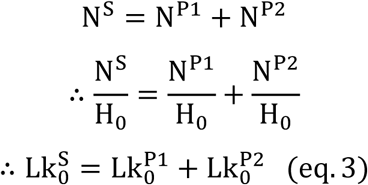

Subtracting eq. 3 from eq. 1 and rearranging we see that the linkage change of the reaction (ΔLk) can be measured as the change in linkage difference between the substrate and the products:

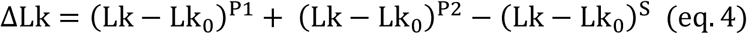

Our strategy for measuring the linkage change of Xer recombination at *psi* was therefore to purify a single topoisomer of a circular substrate plasmid containing two *psi* sites, recombine it *in vitro* with purified Xer proteins and then measure Lk – Lk_0_ for the substrate and separate product circles using gel electrophoresis. The following sections describe two different approaches to these experiments; (a) we used a plasmid containing close spaced *psi* sites such that the small circle produced is a single topoisomer, (b) we used a plasmid with two evenly spaced *psi* sites that recombines to produce two identically sized circles.

### Xer recombination occurs with a fixed ΔLk on the close-spaced substrate

In our first strategy, we utilised a single purified topoisomer of pCLOSE (3437 bp) containing two closely-spaced *psi* recombination sites that recombine to produce a catenane of two circles of 3039 bp and 398 bp (Figure 2a). We predicted that due to its small size, a single topoisomer of the 398 bp circle would be energetically most favourable and would be the predominant product. If this is true, the remaining supercoiling will segregate into the large product circle which will also be a single topoisomer if Xer recombination takes place with a fixed ΔLk.

**Figure 2.**
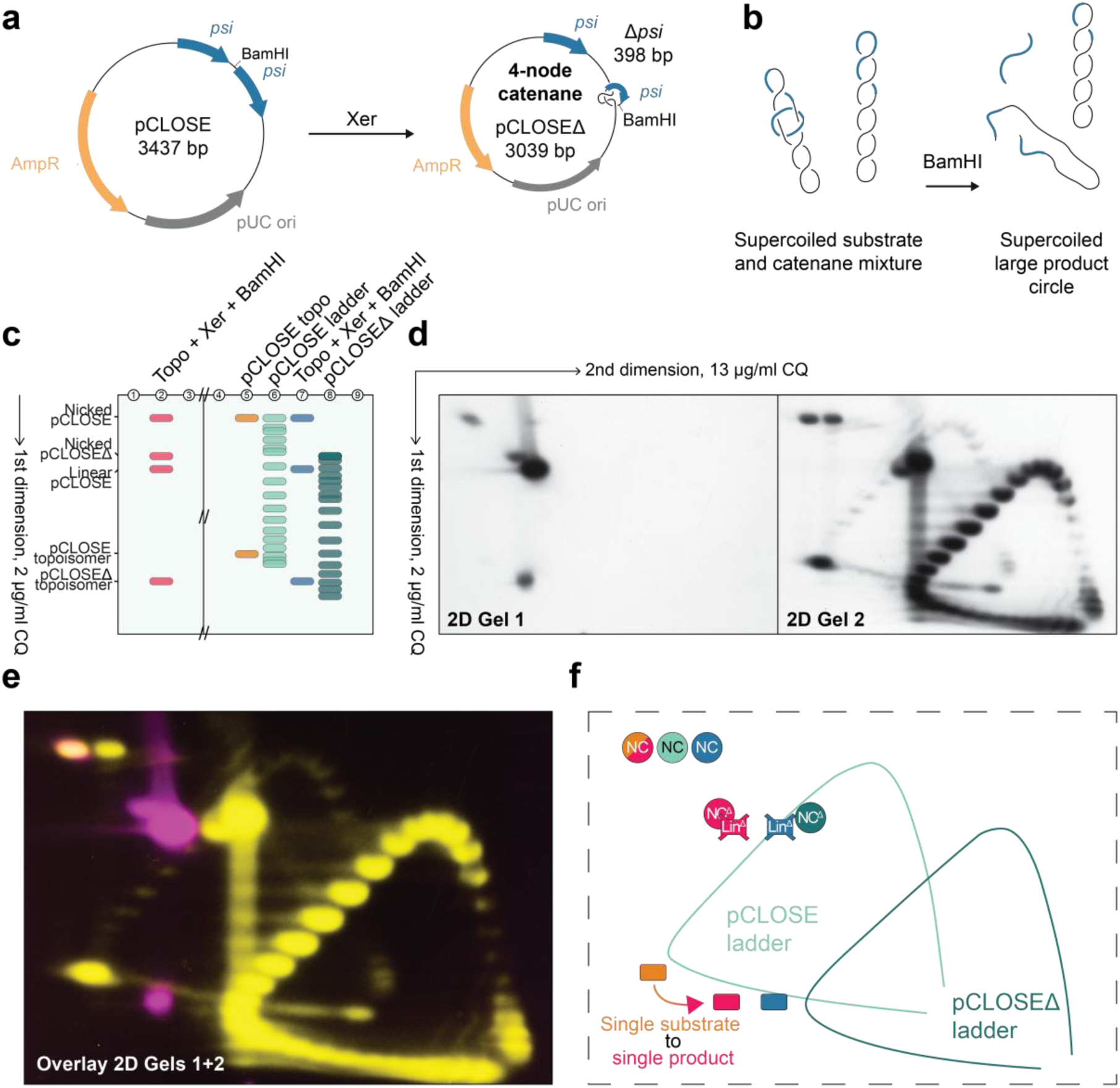
Xer recombination on a single topoisomer of the close-spaced *psi* substrate generates a single product topoisomer. **a**, Maps of the close-spaced *psi* substrate pCLOSE (3437 bp) and its product. Xer recombination of pCLOSE generates a large circle (pCLOSED, 3039 bp) and a small circle (D*psi*, 398 bp) linked together in a 4-node catenane. BamHI restriction endonuclease cleavage sites are marked. **b**, BamHI cleavage of an *in vitro* Xer recombination reaction of pCLOSE linearises the unrecombined substrate and the 398 bp circle in the catenane product, liberating the pCLOSED large circular product for analysis by 2-dimensional gel electrophoresis. **c,** Illustrated migration patterns of the samples present in the first-dimension gel electrophoresis. “Topo” denotes the purified single topoisomer of pCLOSE that was used as the initial substrate for Xer recombination. “Ladders” contain topoisomer mixtures of pCLOSE or pCLOSED. Empty lanes contained no DNA and were included to allow lane 2 and lanes 5-8 to be sliced from the gel separately (cut denoted by the double slashes), rotated through 90° and cast into two new gels before electrophoresis in the second dimension. **d,** Southern blot images after 2-dimensional electrophoresis of lanes 1-3 (2D Gel 1) and lanes 4-9 (2D Gel 2). Gels contained 2 mg/ml chloroquine in the first dimension and 13 mg/ml chloroquine in the second dimension. **e,** Blot images were false-coloured (Gel 1 magenta, Gel 2 yellow) and overlaid with the nicked circle of pCLOSE in lane 2 superimposed on the same species in lane 5. Orange areas correspond to overlapping signal intensity. **f,** Illustration of the DNA species present in (e). The colours used for the different lanes are the same as those used in (c). Abbreviations; NC – nicked circular, Lin – linearised, Topo – topoisomer, D - recombination product.

To test this hypothesis, we purified a single topoisomer of pCLOSE and recombined it *in vitro* with PepA, XerC and XerD. We then used BamHI to cleave the small circle and any unreacted pCLOSE, leaving the large product circle intact (Figure 2b). The BamHI-cleaved Xer-recombined sample was subjected to 2-dimesional gel electrophoresis on its own (Figure 2c, lanes 1-3; Figure 2d, Gel 1), or with the untreated pCLOSE substrate topoisomer, topoisomer reference ladders for the substrate (pCLOSE), and the large product circle (pCLOSEΔ) loaded in adjacent lanes (Figure 2c, lanes 4-9; Figure 2d, Gel 2). All samples were electrophoresed on a single agarose gel containing 2 μg/ml chloroquine in the first dimension (Figure 2c). The two groups of lanes were then separated, rotated by 90°, cast into separate new gels, and subjected to electrophoresis in the second dimension with 13 μg/ml chloroquine (Figure 2d). The DNA species present in each gel were visualised by Southern blotting and hybridisation with ^32^P-labelled pCLOSE DNA.

To aid understanding of the 2-dimensional blots, the layout of the first dimension, showing the loading order and predicted migration patterns of the DNA species in each sample, is shown in Figure 2c. Superposition of the 2-dimensional gel images (Figure 2e), overlaying lane 2 of Gel 1 with lane 5 of Gel 2 allows us to unambiguously assign all of the observed bands (Figure 2f). The most abundant species in the BamHI-cleaved Xer-treated sample is unrecombined pCLOSE that has been linearised with BamHI (Figure 2d, Gel 1). The next most abundant species are the single supercoiled topoisomer of the large product circle (pCLOSEΔ) and its nicked form produced by random strand breakage in this product during sample preparation. The BamHI digestion has not gone to completion so a small amount of the nicked pCLOSE substrate and a very faint band for the unreacted supercoiled pCLOSE topoisomer are also visible on the blot (Figure 2d, Blot 1 and Supplementary Figure S1a). Note that only covalently closed circular molecules change mobility with different concentrations of chloroquine so nicked pCLOSE, nicked pCLOSEΔ, and linear pCLOSE lie on the diagonal of the gel while covalently closed circular topoisomers generally lie off this diagonal. The second 2D gel (Figure 2d, Gel 2) shows two arcs for the topoisomer reference ladders, one for pCLOSE and one for pCLOSEΔ. The nicked (open circular) forms of these plasmids migrate on the diagonals, at approximately the same positions as the slowest topoisomers in both dimensions. The slowest migrating topoisomer in the second dimension (13 μg/ml chloroquine) is more negatively supercoiled than the slowest topoisomer in the first dimension (2 μg/ml chloroquine), so topoisomers become more negatively supercoiled in an anticlockwise direction around the ladders. Any topoisomers in the ladder that become nicked between electrophoresis in the first and second dimensions form a “tail”, streaking down from the nicked band, that is particularly noticeable for the more heavily loaded pCLOSEΔ topoisomer ladder. The position of this tail confirms the identification of the nicked species and the direction in which the gel was run.

The purified substrate topoisomer can be seen as a relatively heavily loaded band to the left of the arc of the pCLOSE topoisomer ladder in gel 2, near to the turning point of the ladder in the second dimension (Figure 2d, 2D Gel 2, lanes 5 and 6). The large product circle in the BamHI-cleaved Xer-treated sample can be seen faintly just to the left of the pCLOSEΔ topoisomer ladder, close to the turning point of this ladder in the second dimension (Figure 2d, 2D Gel 2, lanes 7 and 8; Supplementary Figure S1c for emphasis). Its position is confirmed by overlaying the 2-dimensional blot images (Figure 2e). The overlay clearly shows that a single topoisomer of the substrate is converted to a single topoisomer of the large product circle that migrates faster than the original substrate in both dimensions (Figure 2e,f).

We conclude from these results that Xer recombination takes place with a fixed linkage change (ΔLk) and supercoiling segregates in a fixed pattern into the two product circles derived from pCLOSE. However, we are unable to use this result to precisely measure ΔLk of the Xer reaction because: 1) it proved difficult to count all of the topoisomers in the marker ladders and determine the exact corresponding positions of the substrate and product band (see Discussion) and 2) Lk-Lk_0_ needs to be measured for the substrate and both product circles (3039 bp and 398 bp) in the same conditions to determine ΔLk, and it would not be easy to measure Lk-Lk_0_ for the small circle in the exact same conditions used for either dimension of these gels.

### Measurement of Lk-Lk_0_ on a closely-spaced Xer substrate

In the next experiment, the electrophoresis conditions were modified so that we could count the topoisomers in the marker ladders more readily. Furthermore, chloroquine was removed from the second dimension so that we could measure Lk-Lk_0_ of the substrate and large product circle in the absence of intercalator. Measurement of Lk-Lk_0_ of the small circle product in the absence of intercalator would then allow determination of ΔLk for the Xer reaction.

Four successive topoisomers of pCLOSE (topoisomers A-D) were purified from natively supercoiled plasmid DNA by electrophoresis in the presence of chloroquine. These four topoisomers were recombined *in vitro* using purified XerC, XerD and PepA, then cleaved with BamHI and separated by 2-dimensional gel electrophoresis with 1.25 μg/ml chloroquine in the first dimension and without chloroquine in the second dimension. Untreated topoisomers A-D and BamHI-cleaved Xer-recombined topoisomers A-D were run in separate lanes separated by a marker lane containing mixed topoisomers of pCLOSE and pCLOSEΔ (Figure 3b). The DNA was transferred to a nylon membrane and detected using Southern hybridisation with ^32^P-labelled pCLOSE. As before, we illustrate the loading order and deduced migration of the gel in the first dimension (Figure 3a), to aid understanding of the results shown in the 2-dimensional blot.

**Figure 3.**
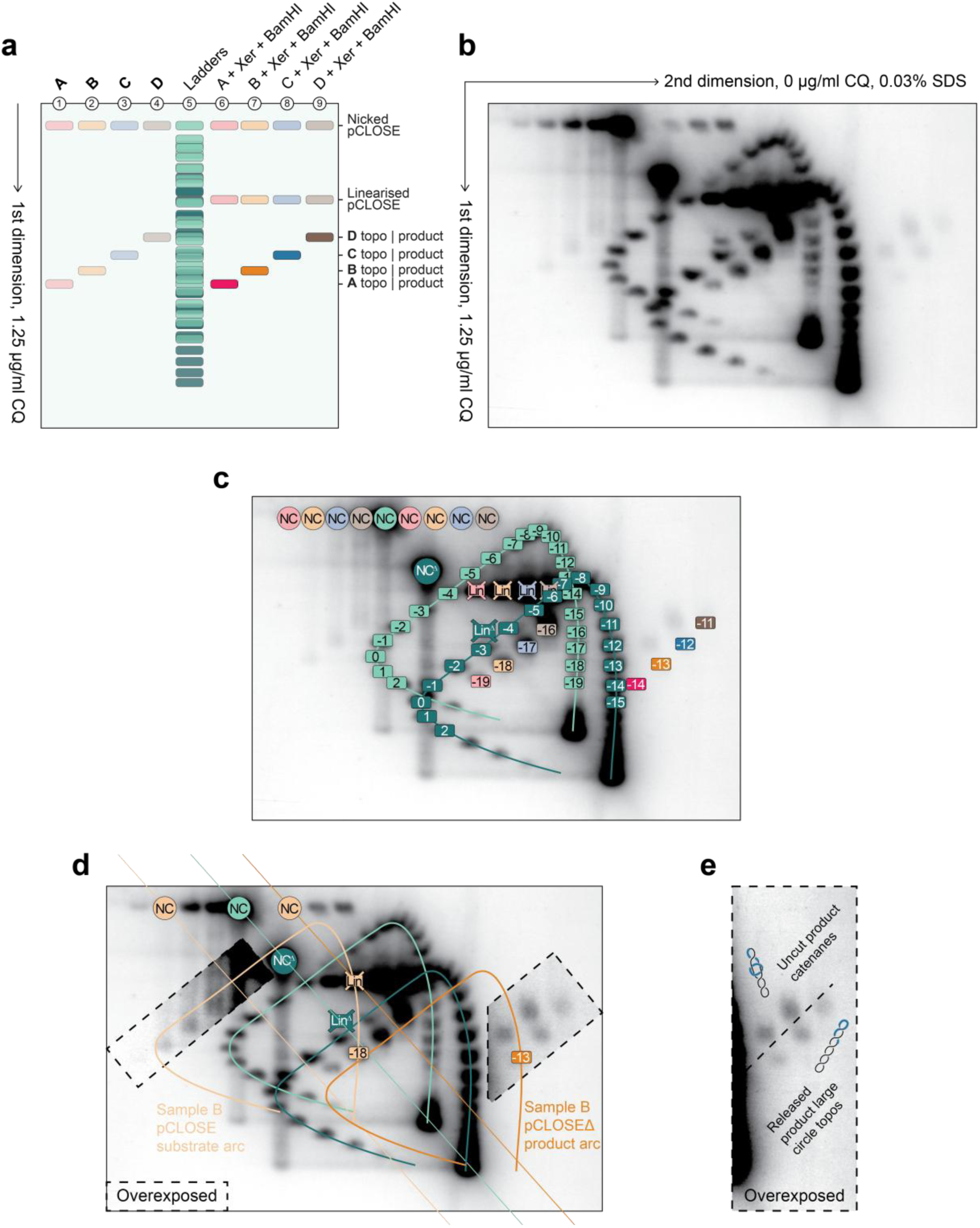
Lk-Lk_0_ of substrate and large product of Xer recombination of pCLOSE. **a**, illustrated mobilities of the samples in the first dimension of gel electrophoresis. Samples A-D are consecutive purified topoisomers of pCLOSE. Topoisomer ladders are mixed populations of pCLOSE (substrate, light-green) and pCLOSED (product, dark-green) topoisomers, loaded together in a single well. Lanes 1-4 contain untreated topoisomers A-D, Lanes 6-9 contain the same topoisomers reacted with Xer and cleaved with BamHI. Species derived from topoisomers A/B/C/D are coloured Red/Orange/Blue/Brown respectively. Light-shaded colours correspond to substrates, while dark-shaded colours correspond to recombination products. pCLOSE and pCLOSED species derived from the ladder lane are coloured light and dark green respectively. **b,** Southern blot of two-dimensional gel loaded as shown in (a). 1.2% agarose TAE gel, containing 1.25 mg/ml chloroquine in the first dimension, then no chloroquine and 0.03% SDS in the second dimension. An overexposed image of this blot is presented in **Supplementary Figure S2a**. **c,** Annotated diagram of the two-dimensional gel blot as shown in (b). Colours are the same as those in (a). Abbreviations; NC, nicked-circular substrate; NCD, nicked-circular pCLOSED; Lin, linear substrate; LinD, linear pCLOSED. Topoisomers are numbered starting from the most relaxed topoisomer in the second dimension. Isolated Xer substrate topoisomers and their pCLOSED products are numbered by comparison to the appropriate topoisomer ladders, as shown schematically in **Supplementary Figure S2b**. **d,** Confirmation that substrate topoisomer B migrates on the arc of pCLOSE topoisomers while its’ cleaved recombination products (-13) migrate on the arc of pCLOSED topoisomers. Topoisomer arcs of pCLOSE and pCLOSED from the ladder lane are shown as light and dark green lines respectively. Nicked pCLOSE (NC), nicked pCLOSED (NCD) and linear pCLOSED (LinD) from the ladder are indicated in the same light/dark green with the gel diagonal drawn as a line through them. The pCLOSE arc is translated 3 (light orange) lanes to the left, superimposing nicked circular pCLOSE bands, to show that topoisomer B (-18) lies on this arc. The pCLOSED arc is translated 2 (dark orange) lanes to the right to show that the species identified as pCLOSED product from B (-13) lies on this arc. Dashed lines indicate areas where an overexposed version of the blot is shown for better visualisation of the low concentration DNA species. See **Supplementary Figure S2c** for similar arc predictions for all topoisomers A-D together, or see **Supplementary Figure S3** for arc predictions shown individually. **e,** Over-exposed section of the blot to the right of the ladder lane showing 4 bands identified as pCLOSED products from topoisomers A-D and 2 bands identified as uncut unrecombined topoisomers C and D.

Two distinct arcs of topoisomers can be seen in the marker lane, one for pCLOSE and one for pCLOSEΔ. Since the second dimension was run with zero chloroquine, the topoisomers become increasingly negatively supercoiled in a clockwise direction around the ladder arcs. Nicked pCLOSE, nicked pCLOSEΔ, linear pCLOSEΔ, and the most negatively supercoiled topoisomers of both plasmids run on the diagonal of the marker lane. As before, a vertical smear can be seen streaking down from both nicked plasmids, representing molecules in the topoisomer ladder that have been nicked between the first and second dimensions. The bands in both topoisomer ladders are well separated, and using the exposure shown in Figure 3b together with alternative exposures, we are confident that we can identify and count all topoisomers in both ladders, starting from those with Lk = Lk_0_, the slowest topoisomers in the second dimension (Figure 3c).

Untreated topoisomers A, B, C, and D were loaded in the four lanes to the left of the pCLOSE marker ladder. Their nicked forms can be seen adjacent to the nicked pCLOSE band in the marker, and 4 bands corresponding to these topoisomers that became nicked between the first and second dimensions can be seen vertically below each nicked species. The purified topoisomers increase in concentration from A through to D, and the band intensities correspond to these concentrations so that topoisomer A that became nicked between the two dimensions can only be seen in a longer exposure (Figure 3d). The supercoiled topoisomers A-D can be seen horizontally in line with their counterparts that were nicked between first and second dimensions (Supplementary Figure S2b), and correspond to topoisomers -19, -18, -17 and -16 in the pCLOSE topoisomer ladder (Figure 3c).

BamHI-cleaved Xer-recombined topoisomers A-D were loaded to the right of the marker lane. Xer recombination has not been very efficient and the most intense band in each lane is the BamHI-cleaved linearised substrate band (“lin” in Figure 3c). BamHI cleavage has been incomplete and consequently nicked pCLOSE substrate is clearly visible for all but topoisomer A (“NC” in Figure 3c). Single topoisomers of the large product circle are clearly seen for all four topoisomers, to the right of the pCLOSEΔ ladder (Figure 3b,c and in the overexposed panel e). These lie exactly as expected, 1, 2, 3, or 4 lanes to the right of the arc of pCLOSEΔ topoisomer markers (Supplementary Figure S2, S3). Counting round the pCLOSEΔ marker ladder from the slowest topoisomer in the second dimension (marked topoisomer “0”), these four products correspond to topoisomers with Lk-Lk_0_ of -14, -13, -12 and -11 respectively. Additional bands near to these product topoisomers (Figure 3b,c,e) do not correspond to positions on the pCLOSEΔ topoisomer arc, we assign these bands as supercoiled unrecombined pCLOSE substrate topoisomers that have not been cleaved by BamHI.

These uncleaved topoisomer bands are most clearly visible for the higher concentration topoisomers C and D, with the predicted position for topoisomer B obscured by the overlapping pCLOSEΔ topoisomer ladder. To illustrate this, we have mapped the 2D arc trajectories for each of the ladders onto the purified substrate topoisomers lanes for topoisomer B (Figure 3d), for all 4 topoisomers together (Supplementary Figure S2c), or for each topoisomer individually (Supplementary Figure S3a-e).

Our conclusion from these results is that pCLOSE topoisomer -19 recombines to form a catenane in which the large circle is topoisomer -14, pCLOSE topoisomer -18 yields catenanes containing pCLOSEΔ topoisomer -15, and so on. Each successive substrate topoisomer yields catenated products containing large (pCLOSEΔ) circles with Lk-Lk_0_ 5 higher than the original substrate. However, to ascertain the overall ΔLk for the Xer reaction of pCLOSE, we must determine Lk-Lk_0_ for the 398 bp circle product, as described in the next section.

### Measuring Lk-Lk_0_ of the 398 bp circular product from pCLOSE

We used 1-dimensional polyacrylamide gel electrophoresis to determine Lk-Lk_0_ for the small (398 bp) circle produced by Xer recombination of pCLOSE. Natively supercoiled pCLOSE DNA was recombined *in vitro* using purified XerC, XerD and PepA. The reaction products were cleaved with HindIII to linearise the unrecombined substrate and the large product circle, releasing the 398 bp circle from the catenane. The HindIII-cleaved products were separated on a polyacrylamide gel in Tris-borate buffer containing 10 mM CaCl_2_ (Figure 4c; lane 5), as calcium ions have been found to improve the separation of small circular topoisomers in polyacrylamide gels ^73,74^. Circular DNA molecules have very low mobility in polyacrylamide gels compared to linear DNA of the same size, so markers for circular and linearised forms of both the pCLOSE substrate (Figure 4c; lanes 1 and 2) and the large pCLOSEΔ product (Figure 4c; lanes 3 and 4) were included on the gel. The HindIII-cleaved recombination reaction sample produced bands that migrated at approximately the correct position for the linearised forms of pCLOSE and pCLOSEΔ, and one major additional band that could be a topoisomer of the 398 bp circle (Figure 4c; lane 5).

**Figure 4.**
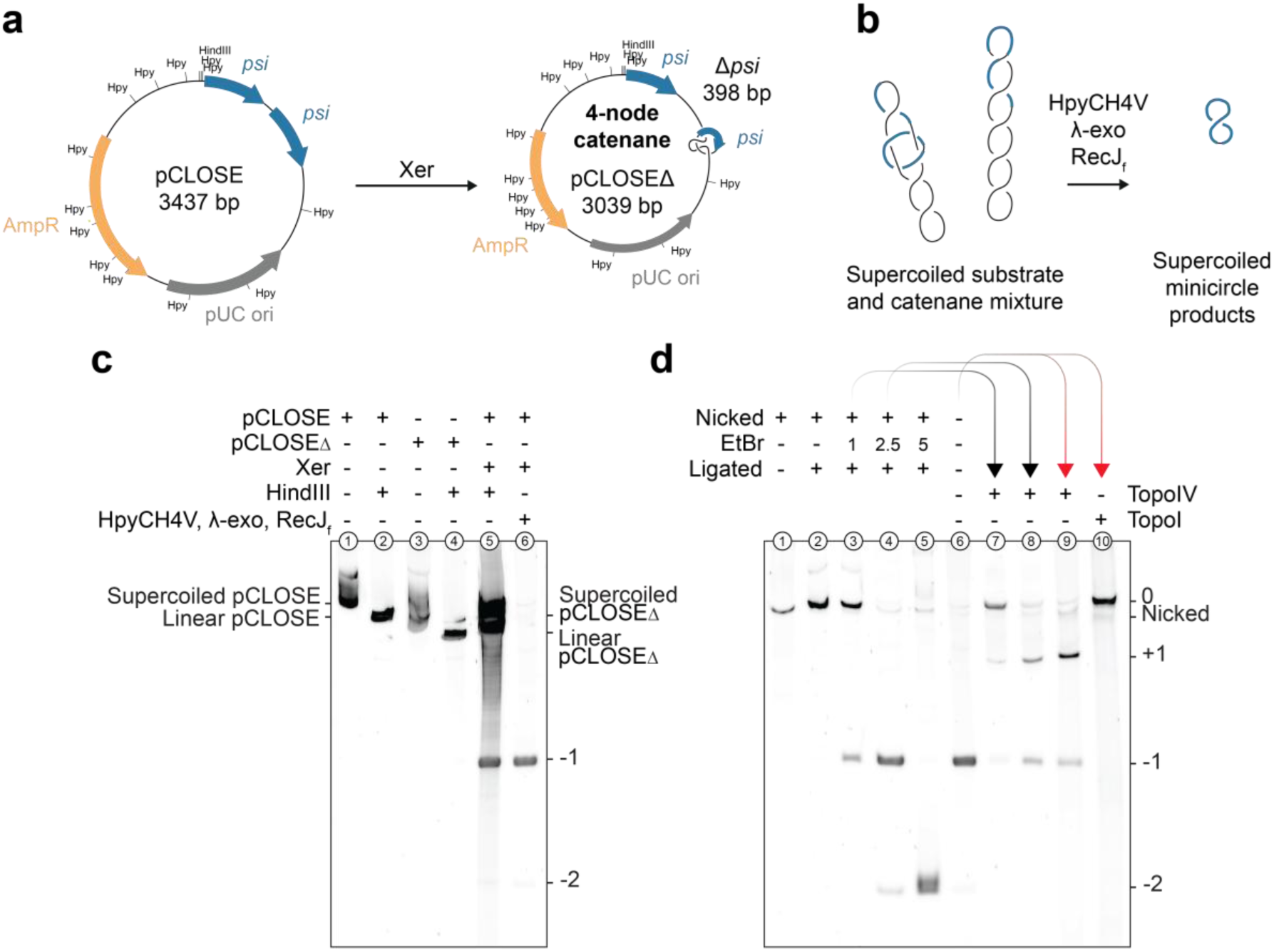
Lk-Lk_0_ of the small product of Xer recombination of pCLOSE. **a**, DNA maps of the pCLOSE Xer recombination substrate and catenated products showing the positions of HindIII and HpyCH4V (Hpy) restriction endonuclease cleavage sites. **b**, The small product circle topoisomers (398 bp) are isolated by removal of the substrate DNA (pCLOSE) and large product circle (pCLOSED). HpyCH4V cleaves pCLOSE and pCLOSED into small linear fragments that are then digested to single nucleotides by the paired actions of l-exonuclease (l-exo) and RecJ_f_, leaving the small product circle intact. **c**, Polyacrylamide gel analysis of the recombination substrate (pCLOSE), the large product circle (pCLOSED), and Xer-recombined pCLOSE, uncut or cleaved with HindIII, or cleaved with HpyCH4V and treated with l-exo and RecJ_f_, as indicated above the lanes. **d**, Polyacrylamide gel comparing 398 bp circle produced by recombination of natively supercoiled pCLOSE (lane 6) to markers produced by reacting the same sample with different enzymes. Treatments are indicated above the lanes. Lane 1, 398 bp circle singly nicked with Nt.BsmAI before removal of pCLOSE and pCLOSED. Lanes 2-5, the nick in the 398 bp circle was resealed with T4 DNA ligase at the indicated concentrations of ethidium bromide to induce different levels of negative supercoiling. Lanes 7-8: Black arrows indicate that the ethidium bromide supercoiled reactions of lanes 3 and 4 were subsequently relaxed with topoisomerase IV. Lanes 9-10: red arrows indicate the natively supercoiled Xer product, present in lane 6, was subsequently relaxed with topoisomerase IV or eukaryotic topoisomerase I.

The linearised substrate, pCLOSEΔ bands, and the background smear of DNA in lane 5 might obscure other topoisomers of the 398 bp circle. We therefore developed a method to deplete the recombination reactions of unrecombined pCLOSE substrate and the pCLOSEΔ product circle, leaving the 398 bp circle intact. The restriction endonuclease HpyCH4V cleaves thirteen times in both pCLOSE and pCLOSEΔ yielding blunt-ended linear fragments ranging in size from 8 bp to 940 bp, but does not cleave the 398 bp small circle product (Figure 4a, Supplementary Figure S4). The linear HpyCH4V products can then be digested to single nucleotides using a combination of λ-exonuclease and the exonuclease RecJ_f_. Neither λ-exonuclease or RecJ_f_ cleaves supercoiled, relaxed, or nicked circular double-stranded DNAs, so the 398 bp small circular product is left intact for further analysis (Figure 4b). Incubation of a pCLOSE Xer reaction with HpyCH4V, λ-exonuclease and RecJ_f_ in a “one pot” reaction for 3 hours resulted in complete removal of the unrecombined pCLOSE, pCLOSEΔ, and the background smear, leaving one major band corresponding to the proposed 398 bp topoisomer in the untreated sample and several other very minor bands (Figure 4c; compare lanes 5 and 6). We show below that the major band corresponds to the -1 topoisomer of the 398 bp circle and that the other minor products are different topoisomers and the nicked form of the 398 bp circle.

First, we determined the migration of linear, nicked and relaxed forms of the 398 bp circle. Treatment of the reaction products with BamHI cuts the 398 bp circle just once and produced a band that migrated much faster than the uncut 398 bp small circle products (Supplementary Figure S5; compare lanes 6-8), demonstrating that none of the species seen in Figure 4c lane 6 are the linearised 398 bp circle. Treatment of the pCLOSE Xer reaction with Nt.BsmAI to nick the 398 bp circle just once, followed by removal of pCLOSE and the large circle using HpyCH4V and the exonucleases, produced a band that migrated slower than the major 398 bp circle product (Figure 4d; compare lane 1 with lane 6). To produce relaxed covalently closed 398 bp circle (with Lk = Lk_0_), the reaction products were nicked with Nt.BsmAI and the nick was then sealed in the absence of intercalator using T4 DNA ligase. After removal of the unreacted plasmid and the large product circle, a band was seen that migrated slightly slower than the nicked 398 bp circle (Figure 4d; lanes 1 and 2). Taken together, these results show that Xer recombination of pCLOSE followed by removal of the large pCLOSEΔ circle (Figure 4c; lane 6, Figure 4d; lane 6) yields a supercoiled topoisomer of the 398 bp circle.

Next, to produce markers for the 398 bp circle with different levels of negative supercoiling, we nicked the reaction products with Nt.BsmAI and re-sealed the nick with T4 ligase in the presence of 1, 2.5 or 5 μg/ml ethidium bromide. Two bands were produced in the presence of 1 μg/ml ethidium bromide, one corresponding to relaxed circle (Lk = Lk_0_) and the other to the untreated 398 bp circle produced by Xer recombination (Figure 4d; compare lanes 2, 3 and 6). No bands of intermediate mobility were seen. This strongly suggests that the small circle product of Xer recombination is the -1 topoisomer. In agreement with this, at 2.5 μg/ml ethidium bromide, the major product was the -1 topoisomer and a small amount of a faster migrating band that we assign as the -2 topoisomer (Figure 4d; lane 4) and at 5 μg/ml ethidium bromide this -2 topoisomer was the predominant product (Figure 4d; lane 5). To confirm these assignments, we treated various samples with *E. coli* topoisomerase IV, which changes linking number in steps of 2, or eukaryotic topoisomerase I which can relax positively or negatively supercoiled DNA in steps of 1. Upon topoisomerase IV treatment of the samples that had been ligated in the presence of 1 μg/ml or 2.5 μg/ml ethidium bromide, the amount of relaxed (Lk = Lk_0_) topoisomer appeared unchanged, while approximately half of the -1 topoisomer was converted into a new species that we propose corresponds to the +1 topoisomer. Since topoisomerase IV changes linkage in steps of 2, the -1 topoisomer cannot be converted to the most relaxed topoisomer. Since +1 and -1 topoisomers have approximately equal energy levels, approximately equal quantities of each are produced at equilibrium. In agreement with these conclusions, when we treated the products of Xer recombination directly with topoisomerase IV and examined the 398 bp circle, slightly more than half of the DNA was converted to the putative +1 topoisomer while the remainder was the -1 topoisomer (Figure 4d; lane 9). In contrast, treatment of the products with eukaryotic topoisomerase I brought about complete conversion to the most relaxed topoisomer (Figure 4d; lane 10).

With the migration of these different topoisomers established on this gel system, we used gel densitometry over five samples of the 398 bp recombination product to quantify the proportions of the different topoisomers present. Across these 5 samples, the -1 topoisomer was on average 97.1% (stdev 1.12%) of the total 398 bp circle, while the -2 topoisomer was 2.9% (stdev 1.12%) (Supplementary Figure S6). From these observations, Lk-Lk_0_ of the vast majority of the small circular product of Xer recombination on natively supercoiled pCLOSE is established as being very close to -1. We discuss why this might be the most favoured topoisomer of the small circle later (see Discussion), but now return to the calculation of ΔLk for Xer recombination of pCLOSE. Using the values for the most highly supercoiled topoisomer A:

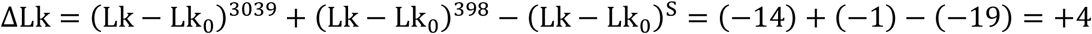

The same result (ΔLk = +4) is obtained with the other 3 topoisomers used (B, C or D).

### The linkage change of Xer recombination observed with an evenly spaced recombination substrate

The close-spaced *psi* sites of pCLOSE ensured that a single topoisomer of the 398 bp circle product was energetically most favourable and therefore the predominant product, making measurement of ΔLk relatively simple. To confirm this measurement, we adopted a second strategy, using a substrate that more closely mimics the natural substrates for Xer recombination, a pseudo-dimeric plasmid (pEVEN) containing equally spaced *psi* sites.

Two monomeric plasmids were constructed by inserting either wild-type *psi* or a *psi* site mutated to introduce a MluI restriction site ^31^ into the micro-cloning vector πAN7 (885 bp) ^66,67^. The two resulting plasmids, pEVENΔ and pEVENΔ*-*MluI (each 1260 bp), were ligated together to create a head-to-tail pseudo-dimeric plasmid (pEVEN; 2520 bp) that contains just one MluI site. Xer recombination of pEVEN produces pEVENΔ and pEVENΔ-MluI linked together in a 4-node catenane (Figure 5a). The two constituent circles of the catenane are identical in size (1260 bp) and identical in sequence apart from the 3 single nucleotide changes made to introduce the MluI site. Treatment of Xer-recombined pEVEN with MluI will linearise any unrecombined pEVEN substrate, and cut one of the two circles from the catenane leaving the other supercoiled (Figure 5b).

**Figure 5.**
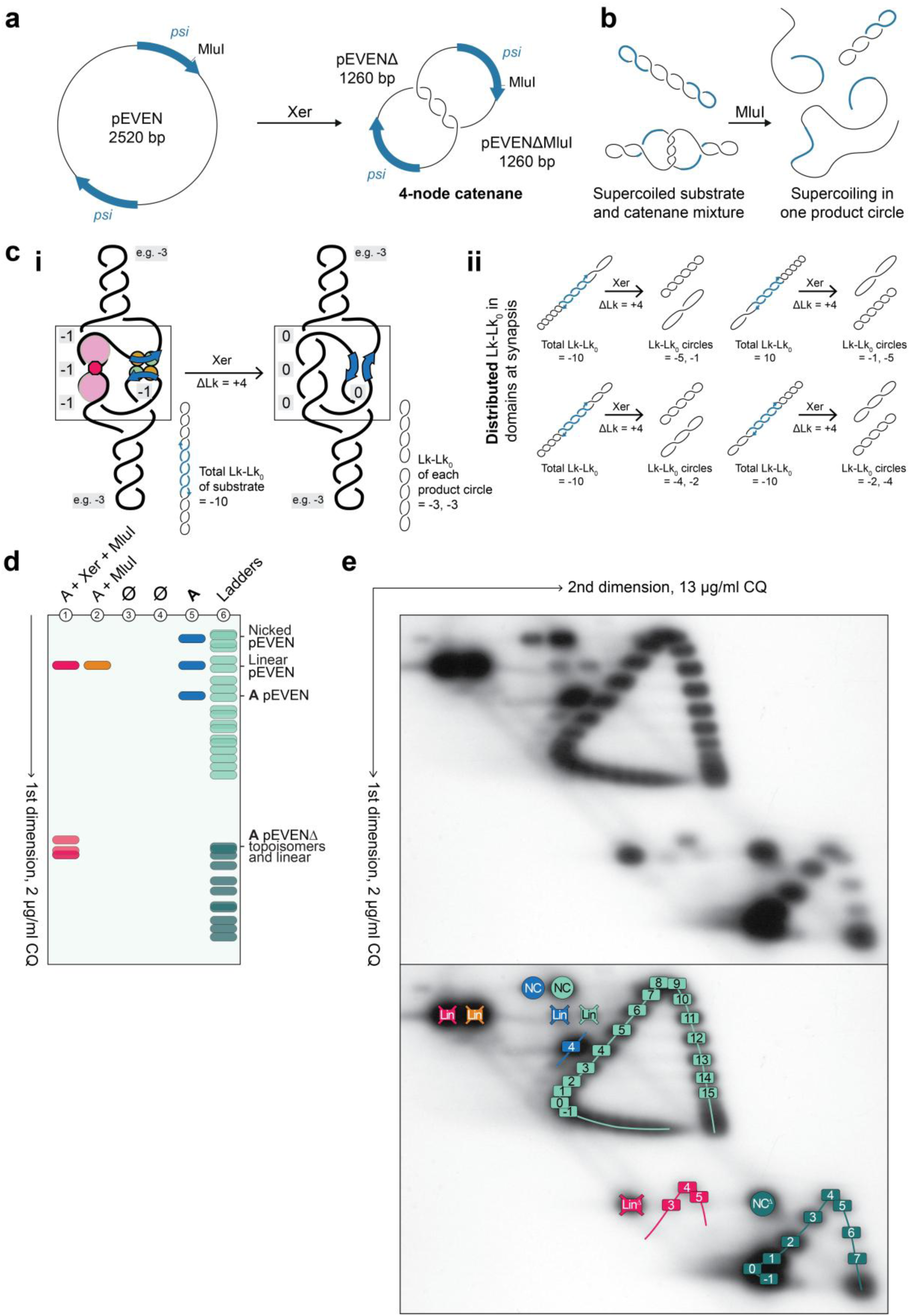
Measurement of linkage change of an evenly spaced substrate. **a**, Maps of the pEVEN recombination substrate and its product, consisting of equal sized catenated circles pEVEND and pEVEND-MluI. The MluI restriction site within one *psi* site is marked. **b**, After Xer recombination of a single topoisomer of pEVEN, one circle of the catenane is linearised by MluI, releasing the other circle for analysis by 2-dimensional gel electrophoresis. Unrecombined pEVEN substrate DNA (2520 bp) is also linearised by MluI. **c**, Partition of supercoiling into the two products of Xer recombination on pEVEN. **i,** A simplified ribbon Illustration showing the -10 topoisomer of pEVEN synapsed to trap 4 negative supercoils constrained by PepA and XerCD binding, with the remaining 6 free negative supercoils partitioned evenly between the two domains that will become the two product circles. Recombination converts 4 negative supercoiling crossings to catenation crossings that do not contribute to Lk of either product circle (0). Symbols: PepA, light-pink circles; ArcA, hot-pink hexagon; XerCD, gold and green circles; recombination core sites, blue arrows. **ii**, Supercoiling can be segregated non-equally into the two domains that will become the two recombination products. The diagram shows alternative ways the supercoils can partition into pEVEND and pEVEND-MluI. Assuming that DLk = +4, pEVEN topoisomer -10 will produce two circles whose total Lk-Lk_0_ will always sum to -6 as shown. **d**, Gel loading order and illustrated mobilities of the samples in the first dimension of gel electrophoresis. Lane 1, pEVEN isolated topoisomer A recombined with Xer, then cleaved with MluI; Lane 2, unrecombined pEVEN topoisomer A cleaved with MluI; Lane 5, untreated pEVEN topoisomer A; Lane 6 (Ladders), mixed topoisomer populations of the pEVEN substrate (light-green) and pEVEND product circle (dark-green). ∅ indicates empty lanes included to separate pEVEND recombination products from the pEVEND ladder after 2-dimensional gel electrophoresis. **e**, Southern blot of 2-dimensional gel loaded as shown in (d), 1.2% agarose TAE gel containing 2 mg/ml chloroquine in the first dimension and 13 mg/ml chloroquine in the second dimension. Upper panel, raw blot image; Lower panel, annotated image of the same gel. Colours used are the same as those in panel (d). Abbreviations: NC, nicked-circular pEVEN; NCD, nicked-circular pEVEND; Lin, linear pEVEN; LinD, linear pEVEND-MluI. Numbers correspond to topoisomer species counted clockwise starting from the most-relaxed topoisomer in the **second dimension** of electrophoresis. The isolated Xer substrate topoisomer A and its pEVEND recombination products are numbered by comparison to the ladders.

The large size of both pEVEN recombination products, compared to the 398 bp product from pCLOSE, reduces the energetic barrier to random segregation of supercoils into the two products during recombination, yielding a distribution of topoisomers for each product circle (Figure 5c). To determine ΔLk on pEVEN, our strategy was therefore to recombine a single purified topoisomer of pEVEN, cleave the reaction products with MluI, and then determine the average linkage difference (Lk-Lk_0_) for the uncleaved product circle using 2D gel electrophoresis. Since both product circles are of identical length and almost identical sequence, both should have identical distributions of topoisomers and we need only examine the linkage difference of one.

In our first experiment using pEVEN, a single topoisomer (topoisomer A) was recombined with Xer proteins *in vitro*, cleaved with MluI, and subject to 2-dimensional agarose gel electrophoresis alongside a range of marker samples. Because the linkage differences of the substrate and product circles are measured in identical conditions on the same gel, we can count the linkage difference relative to Lk_0_ in either dimension, and we do not need to run one dimension in the absence of chloroquine. A mixed topoisomer ladder containing the pEVEN substrate and the pEVENΔ product was generated by relaxing these plasmids with topoisomerase I in the presence of different intercalator concentrations. To produce a linear substrate marker, pEVEN was cleaved with MluI. These samples were then run next to each other on a 2-dimensional agarose gel containing 13 μg/ml chloroquine in the first dimension and 2 μg/ml chloroquine in the second (Figure 5e). As before, we represent the arrangement of the samples and the expected migration of the different species in the first dimension of the gel diagrammatically (Figure 5d).

The topoisomer ladder sample produces two well separated arcs on the 2-dimensional gel: one for pEVEN and one for pEVENΔ, with topoisomers becoming increasingly negatively supercoiled in an anticlockwise direction around the arcs. The pEVEN and pEVENΔ ladders run in distinct parts of the gel, and the topoisomers are well separated and easily countable around both ladders. Nicked circles of pEVEN (NC) and pEVENΔ (NCΔ) are easily identified by their migration on the diagonal, at about the positions of their slowest topoisomers in each dimension. Most of the unreacted pEVEN topoisomer A (Lane 5) runs at the same position as the +4 topoisomer in the pEVEN topoisomer ladder (Lane 6), counting from the topoisomer with Lk=Lk_0_ in the second dimension (13 μg/ml chloroquine). Very small amounts of topoisomers +3, +2 and +1 can be seen in lane 5, presumably due to contamination with neighbouring topoisomers during gel purification. MluI cleavage of Xer-recombined pEVEN topoisomer A (Lane 1) produced linearised unrecombined substrate (Lin) that can be identified using the pEVEN linear marker (Lane 2), as well as linearised pEVENΔ-MluI product (LinΔ), and a distribution of pEVEN topoisomers (red labels in Figure 5e lower panel). Counting from topoisomer 0 in the second dimension, the centre of the product distribution is topoisomer +4, with smaller and approximately equal amounts of topoisomers +3 and +5 (Figure 5e, lower panel; compare red labelled products to green labelled topoisomer ladder). Topoisomer +2 is also just visible, although its abundance is probably exaggerated because it overlaps with a streak on the blot diagonal coming from the linear product on the autoradiogram.

The linkage change ΔLk^pEVEN^ can then be calculated from these results. As discussed above, we assume that on average half the linkage goes into each of the product circles during recombination and therefore that the average linkage differences of pEVENΔ and pEVENΔ-MluI are the same. Therefore

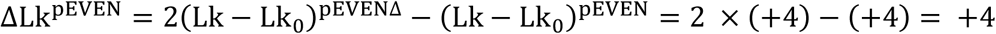

The linkage change can also be calculated by counting the positions of topoisomers on the ladder relative to the most relaxed topoisomers in the first dimension (2 μg/ml chloroquine; Supplementary Figure S7). In the first dimension, two pEVEN topoisomers in the ladder migrated with approximately equal mobility, implying that Lk_0_ is approximately halfway between them at this chloroquine concentration. The two slowest topoisomers therefore have (Lk-Lk_0_) of + and -0.5, and the substrate topoisomer A has (Lk-Lk_0_) = -4.5 at 2 μg/ml chloroquine. In the pEVENΔ topoisomer ladder, there are two topoisomers near the turning point with slightly unequal mobility. Lk_0_ is therefore between these two topoisomers but closer to the slower one and we can approximate (Lk-Lk_0_) = -0.25 for the slowest topoisomer in the ladder which also corresponds to the centre of the product distribution. Putting these numbers into the equation for ΔLk gives:

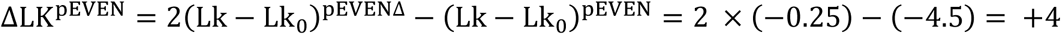

As expected, this is in agreement with the result measured from the turning point (Lk_0_) at 13 μg/ml chloroquine. Thus ΔLk measured with pEVEN is +4, in full agreement with that measured with pCLOSE.

### Linkage segregates as expected into the two pEVEN recombination products

To verify that the linkage segregates equally into the two product circles of pEVEN as expected, we did a second experiment using three consecutive topoisomers of pEVEN (topoisomers A, B, and C where topoisomer A is the same as that used in Figure 5). These topoisomers were recombined with XerC, XerD and PepA, cut with MluI and separated by 2-dimensional electrophoresis in the same conditions as used for Figure 5. Topoisomer ladders were omitted from this gel and the samples were separated at 3-lane intervals so that the pEVENΔ product topoisomer distributions did not overlap. The three untreated substrate topoisomers A, B and C were loaded in adjacent lanes on the same gel to confirm that they were single consecutive topoisomers. As before, the layout of the gel and the deduced migration pattern in the first dimension is shown diagrammatically (Figure 6a). The resulting blot confirms that topoisomers A, B and C are nearly pure and each produces a clear distribution of pEVEN product topoisomers (Figure 6b).

**Figure 6.**
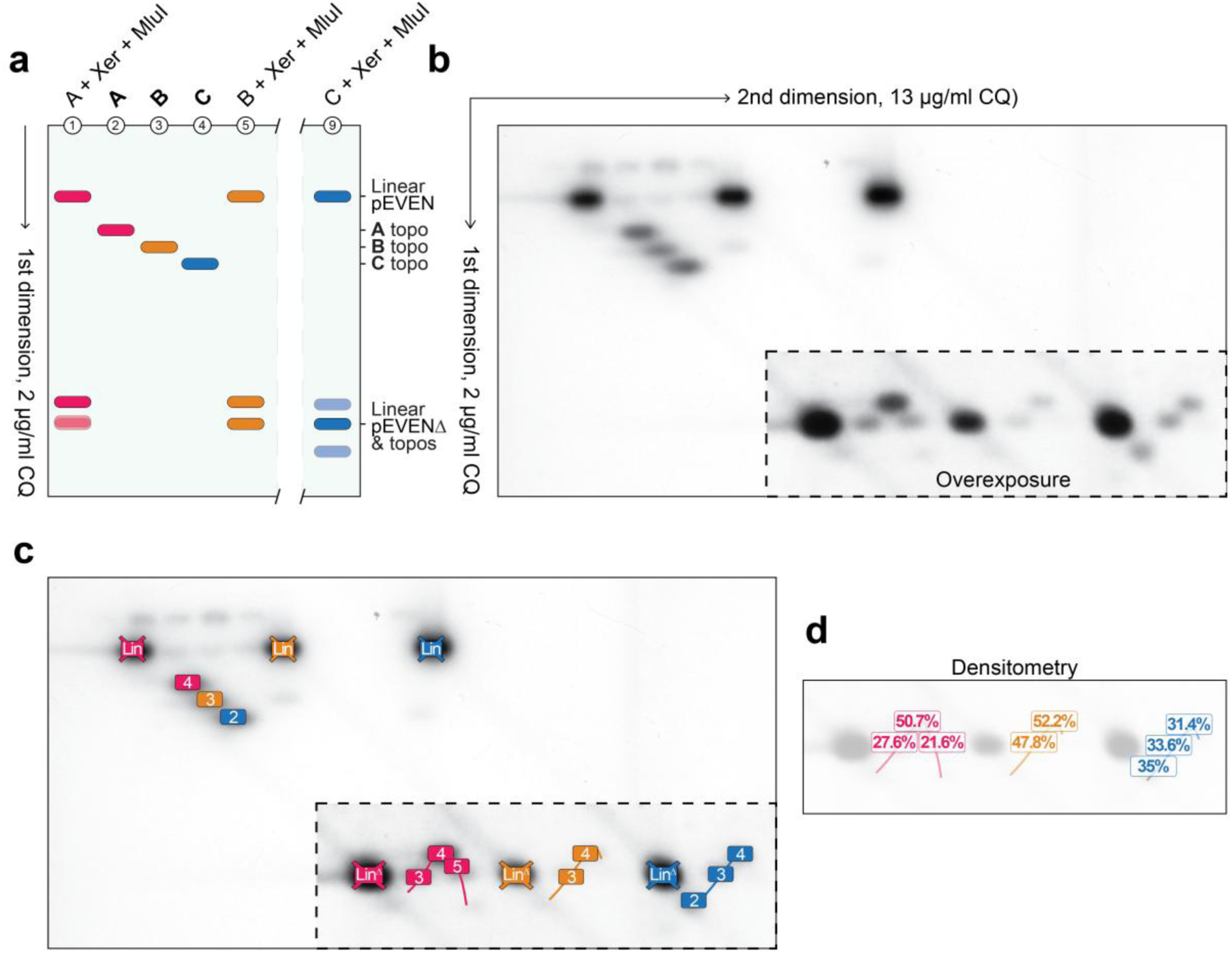
Average Lk-Lk_0_ of pEVENΔ products shift by 0.5 for every unit change of Lk-Lk_0_ in the pEVEN substrate. **a**, Gel loading order and illustrated mobilities of the samples in the first dimension of gel electrophoresis. Lanes 1, 5 and 9; consecutive topoisomers A, B, and C of pEVEN recombined with Xer then cleaved with MluI. Lanes 2, 3, 4; the same topoisomers A, B and C of pEVEN untreated. Three lanes were left between each Xer recombined sample. Topoisomer A is the same DNA as that used in Figure 5. **b**, Southern blot of a 2D gel loaded as shown in (a). 1.2% agarose TAE gel, containing 13 mg/ml chloroquine in the first dimension and 2 mg/ml chloroquine in the second dimension. Dashed lines indicate areas where an overexposed version of the blot is shown for better visualisation of the low concentration DNA species, see: Supplementary Figure S8 for the complete overexposed image. **c**, Annotated diagram of the 2-dimensional gel shown in (b). Colours are the same as those used in (a). Abbreviations; Lin, linear pEVEN; LinD, linear pEVEND-MluI. Numbers correspond to topoisomer species counted clockwise starting from the most-relaxed topoisomer in the second dimension of electrophoresis (deduced from the migration of topoisomer A and its recombination products in Figure 5). **d**, Band densitometry of pEVEN product topoisomers derived from topoisomer A, B and C. Percentages represent the density of a given topoisomer as a fraction of the sum of densities of all pEVEN product topoisomers from the same lane.

Using the assignment of substrate and product linkage differences for topoisomer A deduced from Figure 5, pEVEN topoisomer A was converted by Xer recombination to pEVENΔ topoisomers of +3, +4, and +5, with the +4 species being the most prominent as before (Figure 6b,c). By blot-densitometry these product topoisomers were produced in proportions of 27.6%, 50.7%, and 21.6% respectively (Figure 6d). In turn, pEVEN topoisomer B had (Lk – Lk_0_) of +3 while topoisomer C had (Lk – Lk_0_) of +2, with B yielding almost equal proportions of pEVENΔ topoisomers +3 and +4 (47.8% and 52.2%), while C recombined to produce almost equal proportions of pEVENΔ topoisomers +2, +3, and +4 (35%, 33.6%, and 31.4%) (Figure 6d). Thus, the centre of the product topoisomer distributions shifts by 0.5 linkage units for everyone one unit change of linkage of the substrate, exactly as expected if the linkage is being segregated equally into both product circles and the ΔLk is the same for the reactions on all three topoisomers.

## Discussion

In this study we have determined the precise linking number change (ΔLk) associated with Xer recombination at the plasmid recombination site *psi*. Our results are based on *in vitro* Xer reactions using homogenously purified substrate topoisomers followed by the separation of the recombinant product circles using 2-dimensional gel electrophoresis. We utilised two different *psi* site recombination plasmids, pCLOSE and pEVEN, which contained repeated *psi* sites with either close or equidistant spacing.

Our first experiments with pCLOSE (Figure 2) revealed that Xer recombination between two *psi* sites occurs with a fixed ΔLk; a single topoisomer of pCLOSE produced a catenane containing a single topoisomer of the large product circle. In contrast to other site-specific recombinases which display topoisomerase activity independently from recombination, complicating the measurement of linkage change for these systems ^19,58,64^, there was no sign that the Xer recombinases relaxed the substrate or product DNA in the reaction conditions used in our experiments.

Although this initial experiment clearly demonstrated a fixed linkage in the large recombinant product, it was difficult to determine ΔLk using these gel conditions (Figure 2d). Our best estimate, counting from the most relaxed topoisomers in the marker ladder at 2 μg/ml CQ, is that the pCLOSE substrate had Lk-Lk_0_ of -14 and the product had Lk-Lk_0_ of -10. If Lk-Lk_0_ of the small 398 bp circle product is zero at 2 μg/ml chloroquine, ΔLk for the reaction would be +4, consistent with the later results with pCLOSE and pEVEN discussed below. Although we did not measure the linkage difference for the small circle in the presence of intercalator, the -1 topoisomer produced from pCLOSE (measured in the absence of intercalator; Figure 4) would be expected to be closer to relaxed (Lk-Lk_0_ = 0) in the presence of 2 μg/ml CQ.

Our next experiment, with 1.25 μg/ml chloroquine in the first dimension and without intercalator in the second dimension, gave us a more confident measurement of ΔLk. Under these conditions the substrates, products, and topoisomer ladders were clear and interpretable (Figure 3). We examined Xer reactions using four different purified substrate topoisomers of pCLOSE, with initial linkages varying in steps of 1 (Figure 3c). By comparison against the topoisomer ladder of pCLOSE we measured these substrate topoisomers as having Lk-Lk_0_ values of about -19, -18, -17, and -16 respectively in standard buffer conditions. Following Xer recombination and restriction endonuclease cleavage to remove the small circle, we observed pCLOSEΔ recombination products produced from these pCLOSE topoisomers with Lk-Lk_0_ values of -14, -13, -12, and -11 respectively.

We then analysed the small circle produced by Xer recombination of natively supercoiled pCLOSE containing a range of supercoiling densities overlapping the -19 to -16 topoisomers used above. Cleavage of recombination reactions with a restriction endonuclease together with λ and RecJ_f_ exonuclease removed the substrate and large product circles and allowed characterisation of the small circle by PAGE. Inclusion of Ca^2+^ in gels gave excellent separation of the different topoisomers ^73,74^, and controls using different topoisomerases allowed us to confidently assign the different species observed. This analysis showed that Lk-Lk_0_ of the 398 bp circle produced by Xer recombination on natively supercoiled pCLOSE was almost exclusively -1. Taken together, Lk-Lk_0_ of pCLOSE substrates and the pCLOSEΔ and 398 bp product circles demonstrated that ΔLk of Xer recombination of pCLOSE is +4.

Plasmid pCLOSE was used in these experiments because we predicted that the small size of one of the products (398 bp) would lead to segregation of a fixed number of supercoils into this circle, making it easier to determine ΔLk. The Gibbs free energy of supercoiling (ΔG^sc^) is proportional to the square of (Lk-Lk_0_) and the energetic cost of introducing each supercoil increases disproportionately for circles smaller than about 2000 bp (ΔG^sc^ = K/N (Lk-Lk_0_)^2^) where N = the length of the DNA in bp; K is a constant for N > 2000 bp but for N < 2000 bp, K increases as N decreases) ^75–77^. Making the simplifying assumption that the probability of segregating Lk-Lk_0_ into the two circles is governed by the total free energy of the two product circles, the proportions of different combinations of supercoils in the two products can be predicted using the Boltzmann distribution. Starting from pCLOSE topoisomer -19, assuming that ΔLk is +4 and that Lk_0_ for the small circle is integral, we can predict that the free energy of the product circles is minimised when the 398 bp circle has Lk-Lk_0_ = -1 and the large circle has Lk-Lk_0_ = -14 (Supplementary Table 1). This distribution of supercoiling between the two products is predicted to make up >99% of the total product, in good agreement with our observation that ∼97% of the 398 bp circle is the - 1 topoisomer. According to our calculations, the reduction in free energy from producing a relaxed small circle (Lk-Lk_0_ = 0) would be outweighed by the increase in supercoiling energy of the large circle (Lk-Lk_0_ = -15), while the increased free energy of adding one more negative supercoil to the small circle (Lk-Lk_0_ = -2) far outweighs the reduction in free energy from the less supercoiled large circle (Lk-Lk_0_ = -13). The small amount of 398 bp circle with Lk-Lk_0_ = -2 (∼3%; Figure 4; supplementary figure S6) suggests that in fact the most relaxed topoisomer of the small circle we see on our gels is in fact slightly overwound (i.e. Lk_0_ is not integral) so that formation of the -2 topoisomer is not as unfavourable as predicted by our calculations.

Finally, we measured ΔLk using pEVEN, which recombines to produce two circles of 1260 bp. As predicted from their size, the linkage did not partition into the products of pEVEN in a fixed way; instead a distribution of pEVENΔ topoisomers was observed. We interpreted the results from pEVEN based on the assumption that on average the linkage partitions equally into both product circles due to their identical sizes and near identical sequences. Starting from 3 different substrate topoisomers and using the average linkage difference of the products, we measured ΔLk as +4 for all three substrate topoisomers, exactly the same value obtained with pCLOSE. From the combined results with pCLOSE and pEVEN, we are highly confident that the linkage change during Xer recombination at *psi* is +4.

The linkage change occurring during recombination by other tyrosine recombinases such as λ integrase, Cre and FLP has previously been measured for inversion reactions. Inversion reactions have the advantage that the substrate and products are circles of identical size so that the linkage change can be directly measured as the difference in Lk-Lk_0_ between the substrate and the product on the same gel. The disadvantage of these recombination systems is that they recombine sites after random collision, trapping variable numbers of supercoils between the recombining sites and yielding knotted as well as unknotted inversion products. The linkage change has been measured only for the unknotted inversion products. In these experiments, inversion by λ integrase, Cre and FLP does not proceed with a single fixed linkage change; ΔLk for these reactions is plus or minus 2 ^58,64,78,79^. Positively supercoiled substrates recombine with a linkage change of -2, and negatively supercoiled substrates give ΔLk = +2, ensuring that the reaction is energetically downhill in both cases.

Topological changes that occur during site-specific recombination can be modelled mathematically using tangle equations. This is described briefly here, but see Sumners *et al*. 1995 ^80^ for a more detailed explanation. In tangle analysis, the two DNA crossover sites bound by the recombinases prior to recombination are represented as a ball referred to as the P (for parental) tangle, containing two strings with parallel or antiparallel recombination sites that can be uncrossed, or crossed with a (+) or (-) sign (represented by the (0), (+1) and (-1) tangles respectively; Figure 7a). The two crossover sites divide the substrate circle into two domains that will become the two product circles (for directly repeated sites) when recombination replaces the P tangle with the R (recombinant) tangle. Any tangling between the two domains outside the crossover sites is represented by the O (outside) tangle which can be divided into O_b_, bound by the enzymes and accessory proteins, and O_f_ which contains any remaining free interdomainal crossings to account for the topology of the substrate DNA (Figure 7b).

**Figure 7.**
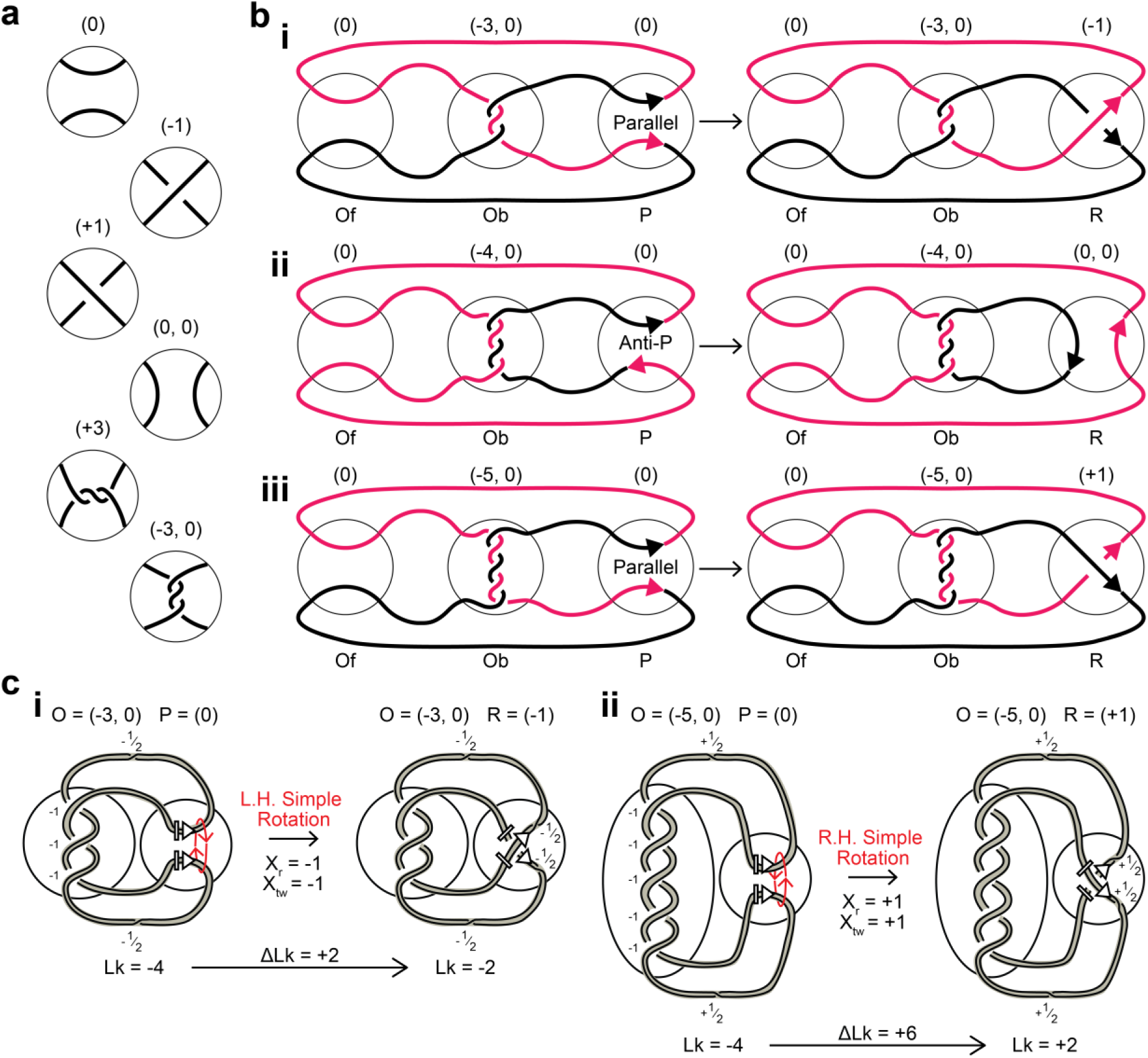
Tangle analysis of Xer recombination at *psi* **a**, Rational tangles. The four trivial tangles and their vector notations (0), (-1), (+1) and (0,0) are shown along with an integral tangle (+3) corresponding to a horizontal row of twists and (-3,0) corresponding to a vertical row of 3 negative twists. **b**, Tangle analysis of site-specific recombination. The substrate is shown as the sum of three tangles: the outside free (Of) tangle not bound by any proteins, the outside bound (Ob) tangle bound by accessory proteins and the parental (P) tangle denoting just the recombination core sites. The core sites in the (P) tangle divide the DNA into two domains coloured black and red. During recombination the P tangle is changed to the recombinant (R) tangle without changing (Of) or (Ob). The three different solutions to the tangle equations for Xer recombination on an unknotted circle are shown with P = (0); Ob = (-3,0), (-4,0) or (-4,0) and R = (-1), (0,0) or (+1) respectively. All yield identical 4-node catenanes with antiparallel *psi* sites as product. **c,** Illustration of predicted linkage change if Xer recombination at *psi* occurred via a simple rotation mechanism. **i,** Left-handed (L.H.) simple rotation with Ob = (-3,0) would yield the observed right-handed 4-node catenane with DLk = +2. **ii**, Right-Handed (R.H.) simple rotation with Ob = (-5, 0) would also yield the observed right-handed 4-node catenane but with DLk = +6.

We initially proposed two possible models for Xer recombination at *psi* to account for the observed production of the right-handed 4-node catenane with antiparallel *psi* sites as the single product topology ^47^. In these models, the crossover sites are initially aligned in parallel and uncrossed (P = (0)) and wrapping of the accessory sequences trap either 3 or 5 negative supercoils (O_b_ = (-3,0) or (-5,0)), with no unbound interdomainal crossings in unknotted circular substrates (O_f_ = (0)). Strand exchange then introduces either one negative (R = (-1)) or one positive (R = (+1)) crossing respectively to produce the observed -4 catenane with antiparallel *psi* sites (Figure 7bi and iii). When crystal structures of Cre and other tyrosine recombinases suggested that these enzymes align the crossover sites in antiparallel ^25,81,82^, we proposed a third model in which the accessory proteins wrap accessory sequences to trap 4 negative supercoils, core sites are aligned in antiparallel and strand exchange produces recombinants also in an antiparallel configuration (O_b_ = (-4,0), P = (0) and R = (0,0); (Figure 7bii)). All three of these models account equally well for the topology of Xer recombination at *psi* on unknotted circles ^31^ and can also account for the products from knotted and catenated DNA substrates ^47,83^. Vazquez *et al.* 2005 ^84^ later demonstrated that under biologically realistic assumptions, these are the only three solutions to the tangle equations for Xer recombination at *psi*, and that all three solutions can represent a single 3-dimensional mechanism seen from three different viewpoints. Note for clarity, there are two possible right-handed 4-node catenanes, with either parallel or antiparallel recombination sites (see: ^47^, Figure 2b). Our previous work demonstrated that the product of Xer recombination at *psi* has antiparallel sites^47^. All three of the tangle solutions discussed above, with either parallel or antiparallel crossover sites in the P tangle, lead to the correct 4-node antiparallel catenane (Fig 7b).

Tangle analysis accounts for the overall topological change during the reaction from unknot to 4-node catenane but does not examine the linkage change of the reaction. To analyse the linkage change we must introduce some further terminology from Stark *et al*. (1989) ^19^. Two parameters X_r_ and X_tw_ are used to summarise the reaction mechanism; X_r_ denotes the local crossings introduced between the two crossover sites by the reaction mechanism, while X_tw_ denotes any twist (positive or negative winding of the two DNA strands around each other) introduced by strand exchange. In the tangle terminology, if P = (0) and the sites are parallel, then X_r_ is the same as the R tangle (assuming R is a simple integer tangle (k)). It can be shown that the sum of X_tw_ and X_r_ gives the change in total linkage of the reaction (ΔLg), which is also equal to the sum of the linkage change of the reaction (ΔLk) and any change in knotting or catenation (ΔKn or ΔCat) ^85^. For reactions between direct repeat sites:

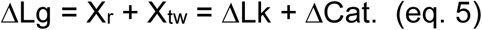

Using eq. 5, and the measured linkage change (ΔLk) we can immediately rule out several possible reaction mechanisms for Xer recombination at *psi*. For instance, ΔLk of recombination by serine recombinases ^19,65^ and X-ray crystal structures of reaction intermediates ^21^ indicate that these enzymes align their sites in parallel and catalyse recombination by a simple rotation mechanism. Simple rotation gives values of X_r_ = +1 and X_tw_ = +1, or X_r_ = -1 and X_tw_ = -1, depending on the direction of rotation (Figure 7c). Using eq. 5 and assuming simple rotation for Xer recombination, the predicted values for the linkage change are ΔLk = +2 for O_b_ = (-3, 0), X_r_ = -1, X_tw_ = -1 (Figure 7ci) or ΔLk = +6 for O_b_ = (-5, 0), X_r_ = +1, X_tw_ = +1 (Figure 7cii), in disagreement with our measured linkage change and thus ruling out simple rotation as the mechanism.

Early models for strand exchange by tyrosine recombinases *via* a HJ intermediate maintained a parallel alignment of sites throughout the reaction and assumed a four-stranded intermediate ^58,59,78,86,87^ (Supplementary Figure 9). Pairwise strand exchange can occur by rotation of the cleaved strands around the uncleaved strands on either side of the four-strand intermediate by either 90° or 270°, giving X_r_ = +1, X_tw_ = -1 or X_r_ = -1, X_tw_ = +1 (ΔLg = 0) for the 90° models, or X_r_ = +1, X_tw_ = +3 or X_r_ = -1, X_tw_ = -3 (ΔLg = +/-4) for the 270° models, all in agreement with the measured linkage change of +/-2 in inversion reactions by λ integrase ^19,58,59^ (Supplementary Figure 9). For Xer recombination at *psi*, our measured linkage change rules out the mechanisms with 270° rotation but is consistent with 90° rotations around the four-strand intermediate. However, the asymmetry of these models, with HJ formation at the start of the reaction not equivalent to the reverse of HJ resolution at the end of the reaction, and later structural results showing a near square planar antiparallel recombination intermediate for tyrosine recombinases, make these reactions mechanisms implausible ^25,58,59,78,81,82,86^. Interestingly, the recent cryo EM structures of IS110/IS621 family RNA bridge-guided recombinases show that they recombine via a parallel-aligned HJ with major grooves aligned towards each other, much like the early models for tyrosine recombinases ^27^. We predict that these RNA-guided recombinases will also catalyse recombination with no net change in linkage (X_r_ = +1, X_tw_ = -1, ΔLg = 0; Supplementary Figure 10).

Symmetric models for tyrosine recombinases have been proposed with the enzyme recognising equivalent chiral structures in the forward and reverse directions and with a square planar HJ intermediate, with sites aligned either parallel or antiparallel ^19,88^(Supplementary Figure 11). In the parallel cases, the first strand exchange produces a parallel stacked HJ that rotates by ∼180° to form an antiparallel HJ, isomerises to cross the other pair of strands, rotates in the other direction to reform a parallel HJ, and finally exchanges the second pair of strands to form the recombinant (Supplementary Figure 11a, c). If the recombinant strands in the initial HJ form a positive crossing, then P = (0), R = (+1) (X_r_ = +1, X_tw_ = -1) (Supplementary Figure 11a). If the initial HJ contains negatively crossed recombinant strands P = (0), R = (-1) (X_r_ = -1, X_tw_ = +1) (Supplementary Figure 11c). Alternatively, the core sites could synapse initially in an antiparallel or near antiparallel conformation (Figure 8a). Strand exchange would produce a HJ that, with only minor rearrangements, can be resolved to the recombinant product with antiparallel sites, with little or no movement of the DNA ends emerging from the complex (Supplementary Figure 11b). This fully antiparallel mechanism, with P = (0) antiparallel and R = (0, 0) antiparallel, does not fit so well into the X_r_ X_tw_ paradigm, but we denote it as X_r_ = (0,0) and Xt_w_ = 0. Taking into account the movement of DNA strands outside the HJ, these three models are predicted to occur with no net change in linkage (ΔLg = 0) (Supplementary Figure 11) and are thus all consistent with our observed ΔLk of Xer recombination, and the actual 3-dimensional reaction mechanism could be anywhere between these fully parallel and fully antiparallel extremes. In fact, if the sites are not exactly co-planar, a given 3-dimensional mechanism will appear parallel from some viewpoints and antiparallel from others, and X_r_ and X_tw_ (but not ΔLg) will change accordingly (Figure 8). One limitation of the tangle model for recombination is that it quantises the crossings by in effect projecting them onto the plane from one chosen viewpoint, while the actual 3-dimensional mechanism may be anywhere between these fully parallel and antiparallel extremes. While evidence of a right-handed crossed conformation of sites during initial synapsis by tyrosine recombinases has been inferred from the bias in product topology ^64^ and by loop closure experiments ^63^ definitive information about the site alignment in Xer recombination at different stages of the reaction awaits further structural studies.

**Figure 8.**
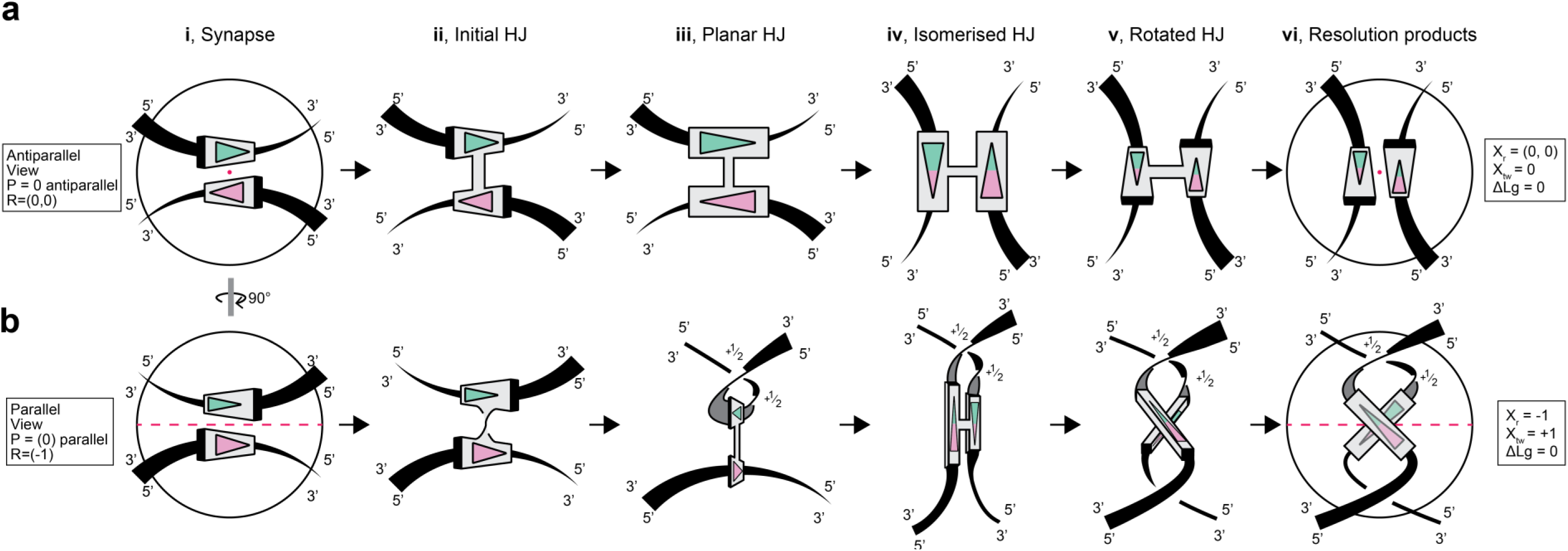
A fully symmetric Holliday junction strand exchange mechanism for tyrosine recombinases. The same mechanism can appear as either parallel or antiparallel from different viewpoints, with different values of P and R tangles, Xr, and Xtw, but with the same total change in linkage ΔLg = 0. **a**, Antiparallel view of strand exchange by tyrosine recombinases. (i) Sites synapse with a right-handed (+) crossing between core sites in 3-dimensional space appear as the antiparallel (0) tangle in this projection. Core sites within which strand exchange takes place are denoted as “dominoes” with green and pink triangles indicating their directions. Black ribbons (linkers) connect the core sites to the boundaries of the tangle ball (circle). Ribbon and domino thickness is used to indicate the 3-dimensional nature of the diagram with thick ends coming out, and thin ends descending below the plane of the paper. The red dot indicates the 2-fold axis of rotational symmetry maintained throughout the mechanism. In subsequent steps of the mechanism, the core sites (dominoes) are cut, rearranged, and re-joined but the ribbon connections to the boundaries of the tangle ball remain unchanged. 5’ and 3’ ends of each ribbon are indicated to keep track of changes in twist during recombination. (ii) Exchange of the first pair of strands produces a HJ. (iii) Clockwise rotation of each site produces a fully planar antiparallel HJ. (iv) Isomerisation of the HJ stacks the recombinant strands and puts the non-recombinant strands into a crossed conformation. (v) Core sites rotate anticlockwise to form a crossed HJ. (vi) Exchange of the second pair of strand produces the fully recombinant product. The core sites in (v) and (vi) have the same right-handed (+) sign crossing as in (ii) and (i). This fully symmetrical model allows the forward and reverse reactions to be structurally equivalent. Overall, the mechanism appears as P = (0) antiparallel, R = (0,0) antiparallel in this projection; X_r_ = (0,0), X_tw_ = 0 and there is no net change in linkage associated with the mechanism (DLg = 0). **b**, The exact same mechanism as in (a) but rotated by 90° around a vertical axis. (i) In this projection, although the core sites form the same right-handed (+) crossing as in (a i), the synapse appears as P = (0) parallel. (ii) First strand exchange produces a HJ with crossed strands that form a left-handed (-) crossing that sets the topology of the final product. (iii) clockwise rotation of both cores forms a planar stacked HJ perpendicular to the plane of the paper. This rotation introduces half a positive twist into each top linker to compensate for the negative crossing introduced between these linkers. (iv) HJ isomerisation swaps crossed and stacked strands, (v) anticlockwise rotation of both cores and (vi) second strand exchange to produce the recombinant product. In the final product, the core site (dominoes) forms a right-handed (+) crossing, but the overall tangle diagram taking the other two crossings into account is (-1). Overall, the mechanism appears as P = (0) parallel, R = (-1) in this projection; X_r_ = -1, X_tw_ = +1 and there is no net change in linkage associated with the mechanism (DLg = 0).

The results obtained for the linkage change during recombination by other tyrosine recombinases (+/-2 for inversion by λ Int, Cre and Flp) are also consistent with the reaction proceeding with no net change of ΔLg. The simplest way to understand the variable ΔLk seen in these reactions is that the crossover sites are aligned antiparallel with one crossing elsewhere in the DNA required to allow this alignment ^64^. The supercoiling associated with this crossing changes sign during recombination and if no further twist is introduced by the strand exchange mechanism (ΔLg = 0), ΔLk will be -2 if the external crossing is positive, and +2 if the external crossing is negative (Supplementary Figure 12). In the tangle notation, O = (+1) or (-1), P = (0) antiparallel and R = (0, 0).

We conclude from our results that Xer recombination proceeds with an extraordinarily precise mechanism. Synapsis and movements of the DNA duplexes are constrained to produce a specific 4-node catenane; while precisely defined movements of the DNA strands during strand exchange lead to a fixed linkage change. This contrasts to other tyrosine recombinases, where recombination occurs after random collision of sites, trapping variable numbers of crossings and giving variable product topologies and linkage change. The +4 linkage change of Xer recombination, which converts four negative supercoils to four more energetically favourable catenation crossings, provides a driving force for the reaction. It seems likely that formation of the interwrapped synaptic complex by accessory proteins and accessory sequences positions the crossover sites of *psi* in a favourable conformation for initiation of strand exchange by XerC, and that after completion of recombination by XerD, the XerC-XerD-crossover site complex dissociates to release the change of linkage into the two catenated product circles, making the reaction essentially irreversible.

Remarkably, evolution has converged upon very similar topological filter mechanisms to enforce intramolecular recombination during the resolution of transposon cointegrates. Binding of serine recombinases such as Tn3 and γδ resolvases to their *res* (recombination sequence) accessory binding sites II and III interwraps the two sites ∼3 times to stimulate strand exchange at *res* core site I ^89^. An analogous architecture is utilised by the tyrosine recombinase transposon resolvase from Tn4430 ^90^. Interestingly, the *psi* topological filter is only one of several mechanisms that have evolved to regulate XerCD activity. Bacterial chromosome dimer resolution by XerCD is driven by FtsK through translocation and alignment of the two *dif* sites of a dimeric chromosome at the divisome, followed by the activation of XerD-first recombination *via* protein-protein contacts between XerCD and the extreme C-terminal γ-domain of FtsK ^1^.

## Supplementary data

Supplementary Data are available at the end of this document.

## Acknowledgements

We thank Martin Boocock and Marshall Stark for their expertise, inspiring discussions, and invaluable comments.

## Author Contributions

Contributions according to CrediT taxonomy in author order: Conceptualization:

S.D.C. and D.J.S; Funding acquisition: D.J.S. and S.D.C.; Formal analysis: S.D.C.; Investigation: J.I.P, A.O.T, and S.D.C.; Methodology: D.J.S. and S.D.C.; Vizualization:

J.I.P. and S.D.C.; Writing – original draft: J.I.P. and S.D.C.; Writing – reviewing & editing: J.I.P., D.J.S. and S.D.C.

---

**Supplementary Figure S1.**
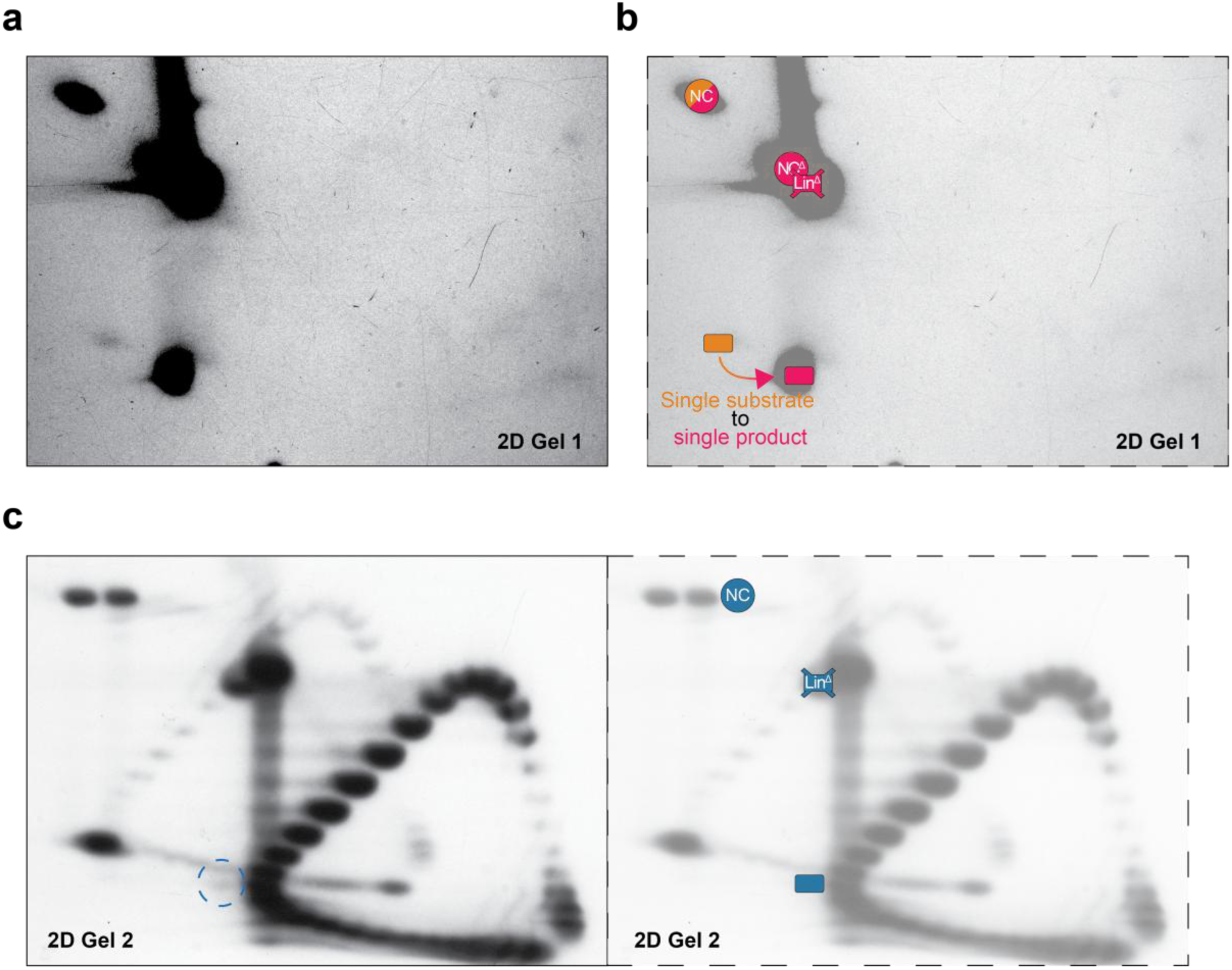
Additional individual annotations of Figure 2d and 2f **a**, Exposure adjusted version of 2-dimensional Figure 2d **Gel 1** revealing the small proportion of unreacted and non-linearised original pCLOSE substrate topoisomer in comparison to the single product topoisomer formed of pCLOSED. **b**, Illustration of the DNA species superimposed on (a) using the same symbols and positioning shown in Figure 2f. Abbreviations; NC – nicked circular, Lin – linearised, Topo – topoisomer, D - recombination product. **c**, Additional annotations of Figure 2d Gel 2. Left, a blue dashed circle marks the location of the low concentration supercoiled recombination product band originally contained within gel lane 7 (Figure 2c). Right, the annotations of Figure 2f for gel lane 7 on the blot. Abbreviations as above.

**Supplementary Figure S2.**
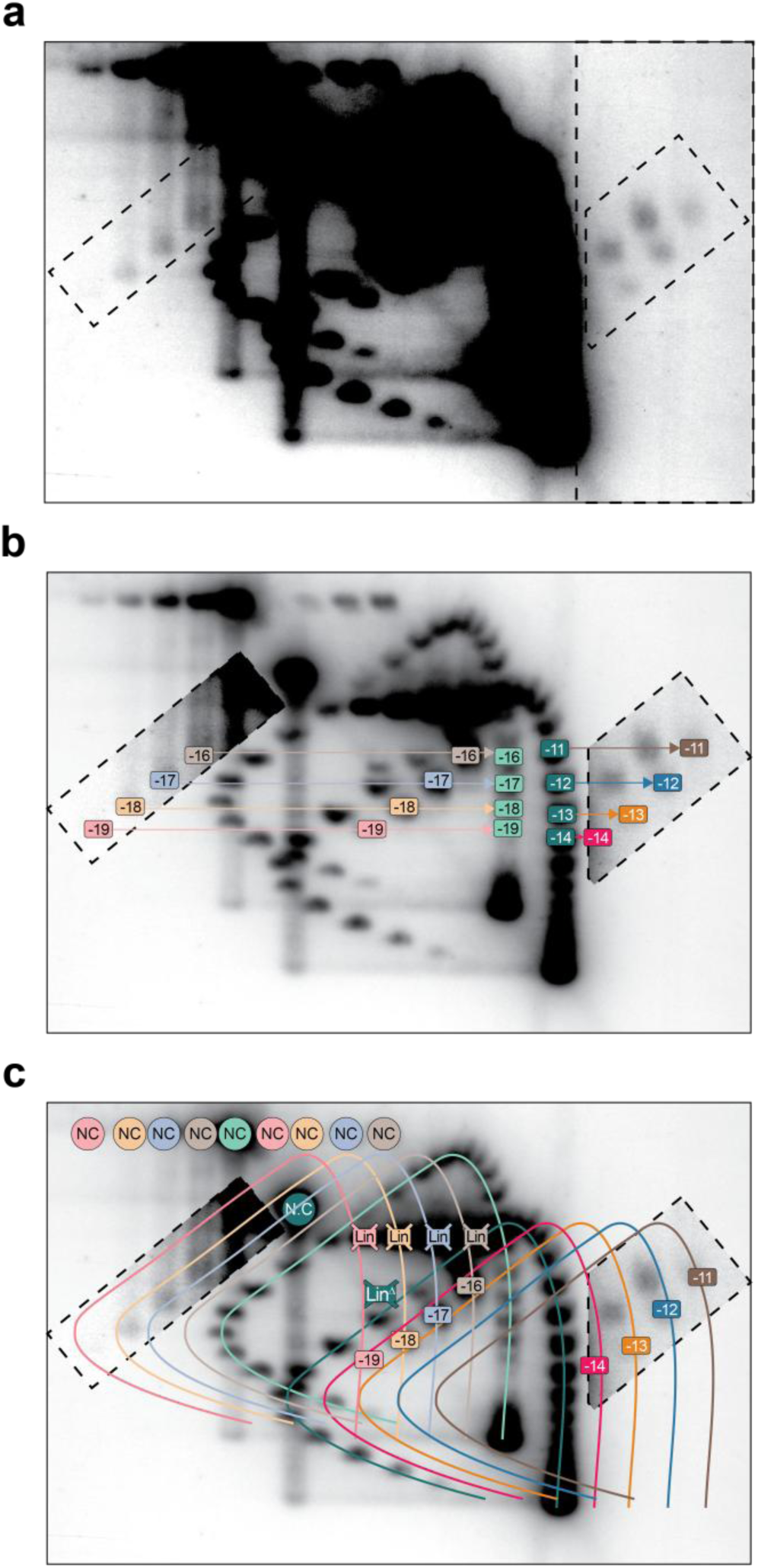
Additional annotations for pCLOSE 2D gel species. **a**, Overexposed version of the blot featured in Figure 3b. Dashed sectors represent areas of overexposure that are superimposed in other subfigures. **b**, Annotation of the isolated pCLOSE substrate topoisomers versus the substrate ladder, and pCLOSED products versus the product ladder. Numbering is the same as in Figure 3c. A small proportion of topoisomers A to D loaded in lanes 1-4 (numbered -16 to -19 in left dashed sector) became nicked during gel processing between the first and second gel dimensions. These species migrate vertically below the bands that had become nicked before gel electrophoresis, and horizontally to the left of corresponding topoisomers loaded in these lanes that remained supercoiled throughout (central bands numbered -16 to -19; coloured light red, orange, blue and brown). The bands in the pCLOSE substrate ladder that these 4 substrate topoisomers correspond to are numbered -16 to -19 in light green. The four pCLOSED product topoisomers loaded in lanes 6-9 are coloured dark red, orange, blue and brown and numbered -14, -13, -12 and -11 in the rightmost dashed sector. The topoisomers they correspond to in the pCLOSED ladder are coloured dark green. Horizontal arrows indicate corresponding bands. **c**, Substrate and product topoisomer arcs corresponding to each sample. The pCLOSE topoisomer arc is shown translated 1, 2, 3, or 4 lanes to the left (light brown, blue, orange or red lines respectively) to indicate where bands corresponding to topoisomers D, C, B, or A should migrate. The pCLOSED topoisomer arc is translated 1, 2, 3, or 4 lanes to the right (dark red, orange, blue or brown lines respectively) to show where bands corresponding to the large circular product of Xer recombination of topoisomers A, B, C and D should migrate. DNA species which do not rest upon the arcs do not correspond to either the circular substrate or circular product.

**Supplementary Figure S3.**
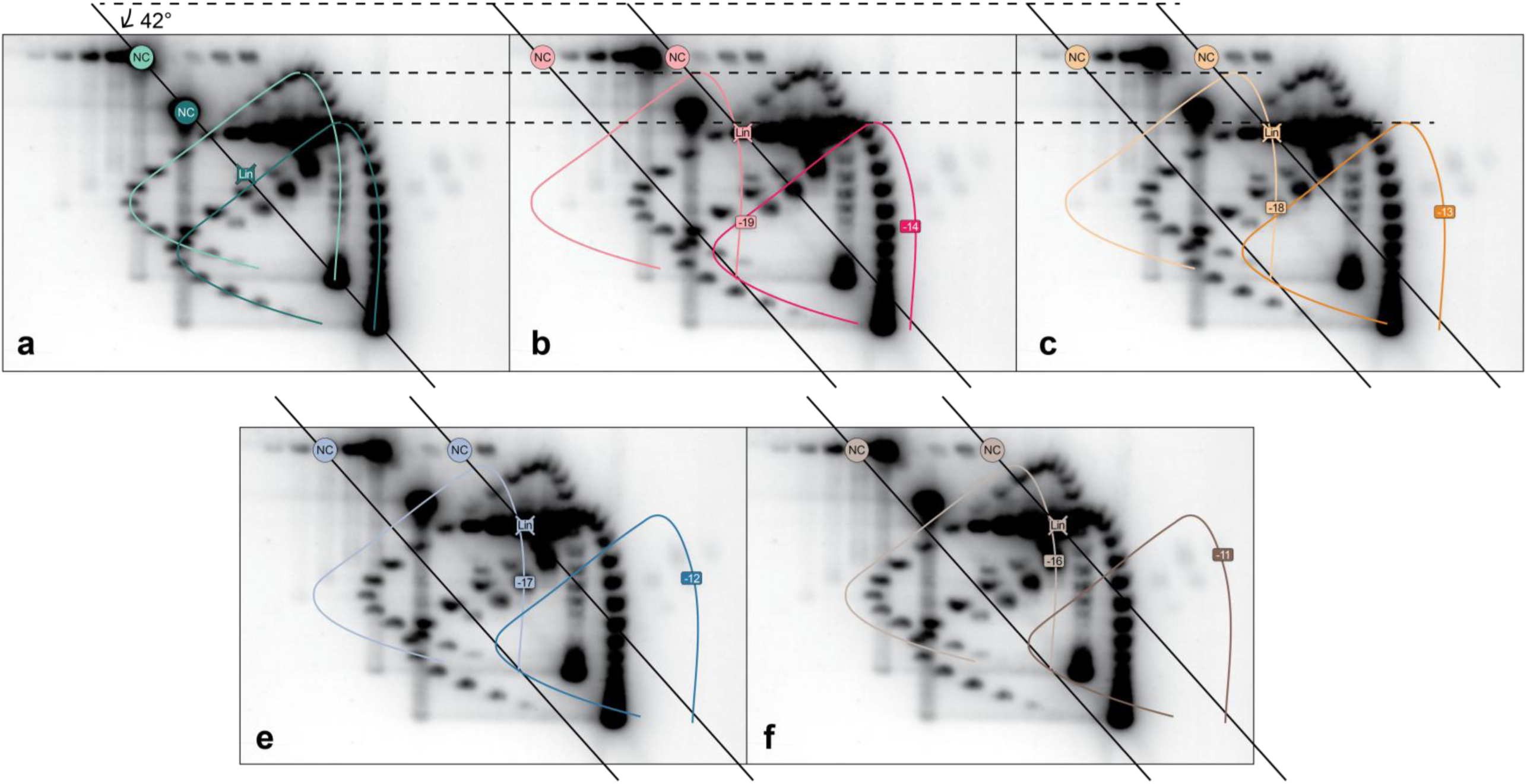
Additional annotations for pCLOSE 2D gel species. Fully annotated substrate and product topoisomer arcs corresponding to the pCLOSE and pCLOSEΔ topoisomer ladders (a), and each isolated pCLOSE topoisomer sample A-D (b-f respectively) coloured as in Supplementary Figure S2. DNA species which do not rest upon the arcs do not correspond to either the circular substrate or circular product of that specific Xer reaction.

**Supplementary Figure S4.**
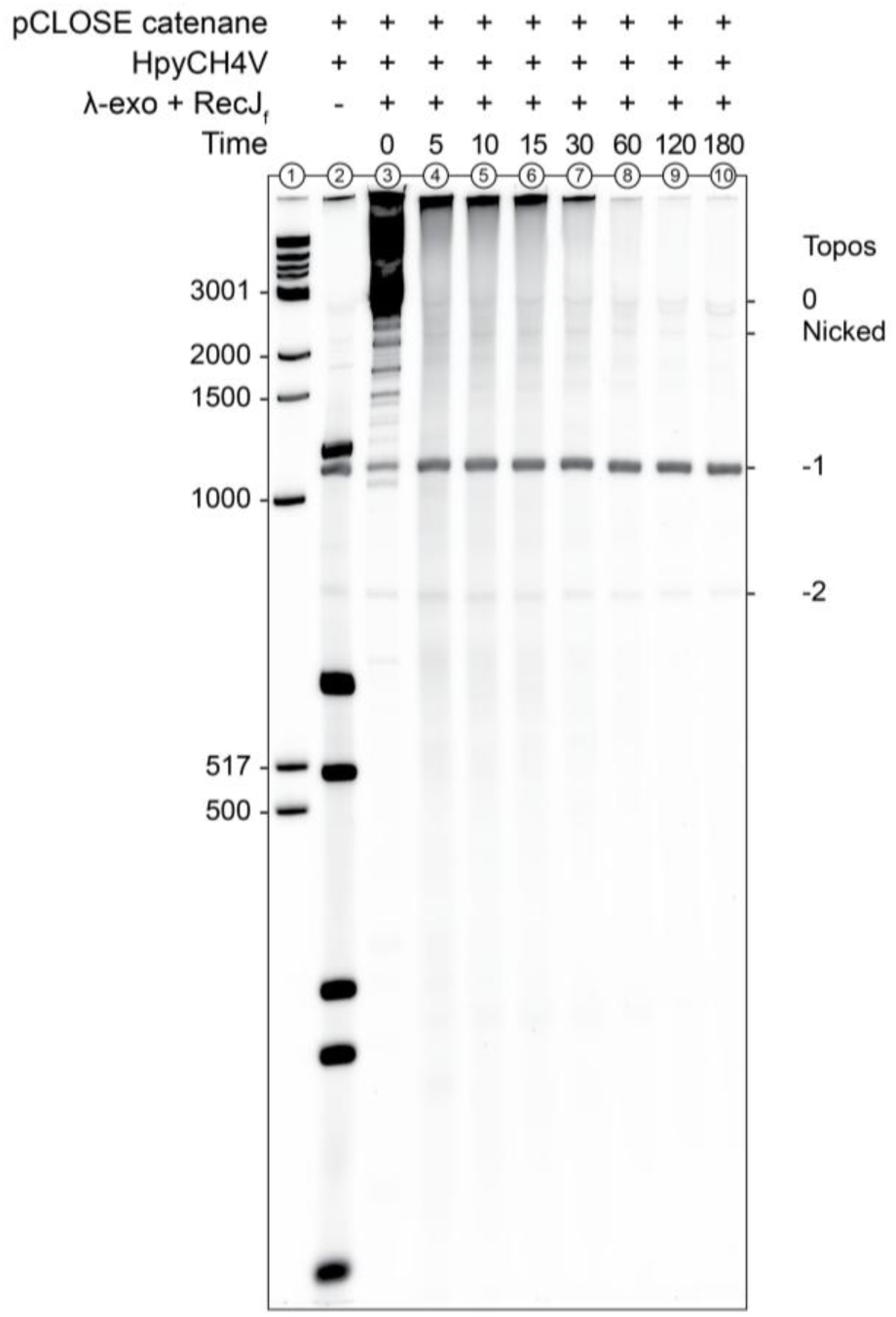
**One-pot endo/exonuclease depletion timecourse of the pCLOSE catenane large circle**. PAGE demonstrates the rapid digestion and disappearance of the pCLOSE catenane large circle (3039 bp) between 0 and 180 minutes when incubated with HpyCH4V restriction endonuclease, λ-exonuclease, and RecJ_f_ exonuclease. Lane 1 contains 1kb DNA Ladder (NEB). Lane 2 contains Xer catenane fragmented with HpyCH4V without exonucleases. Ladder sizes are indicated in bp to the side of the gel image.

**Supplementary Figure S5.**
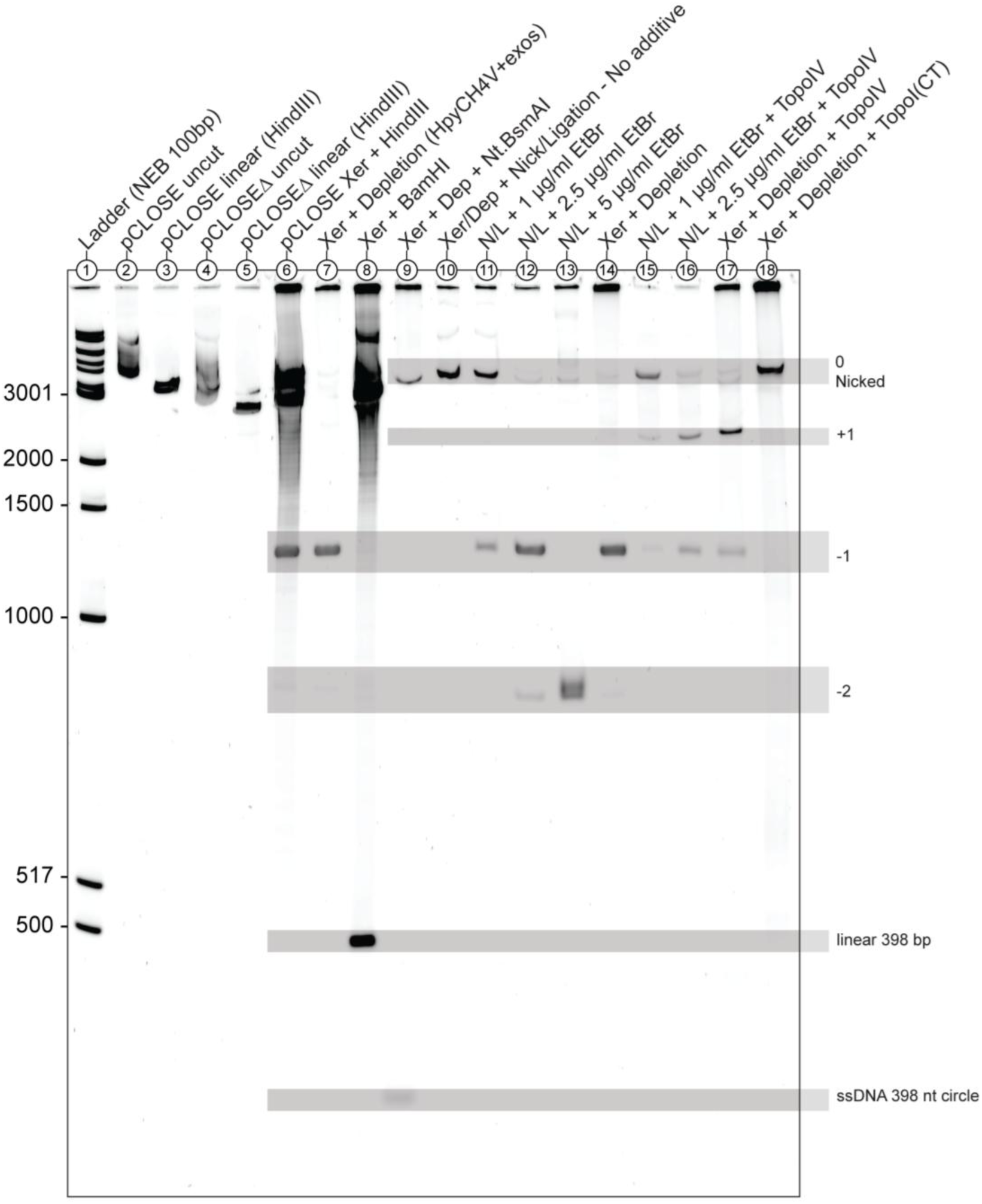
Uncropped version of. Figure 4c-d. 5% PAGE gel containing Tris-borate buffer with 10 mM CaCl_2_, post-stained with SYBR Gold. DNA ladder is New England Biolabs 100 bp (N3231), numbers on the left-hand side mark the major ladder bands in basepairs. Abbreviations; Dep, DNA treated with the depletion reaction of HpyCH4V with exonucleases (exos); Xer, the pCLOSE DNA was recombined with PepA, XerC, and XerD as per methods. N/L, a nick/ligation reaction where nicked pCLOSE catenane was incubated with DNA intercalators before addition of T4 DNA ligase. ssDNA, single-stranded DNA. Nt, nucleotides.

**Supplementary Figure S6.**
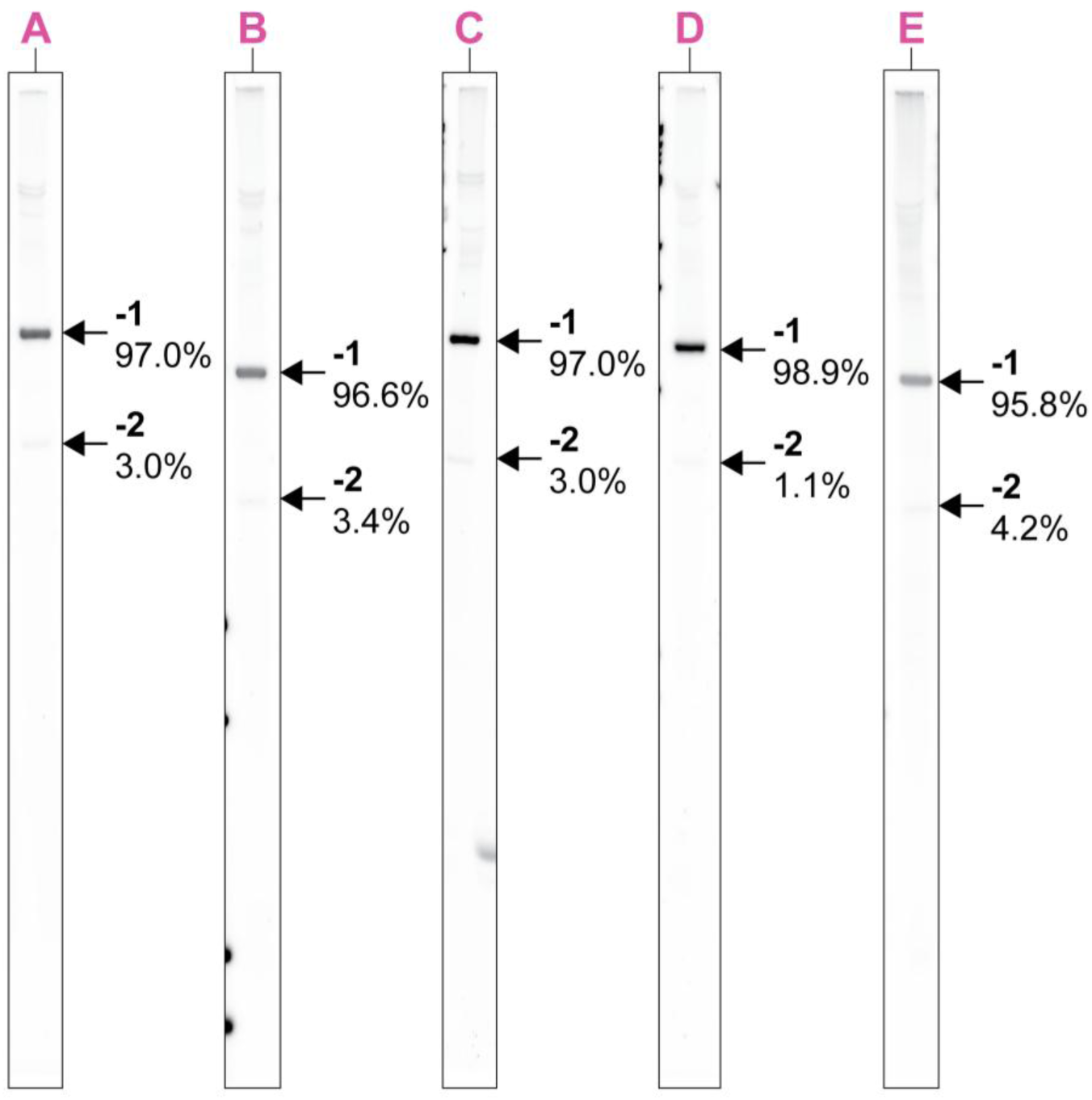
Band densitometry of the pCLOSE small circle recombination products. Each lane (A-E) contains DNA derived from the same pCLOSE Xer recombination reaction which was treated with HpyCH4V and exonucleases for depletion of the large circle product. Each lane is derived from a separate PAGE gel. Lane contents were examined by band densitometry and represented here as percentages of the sum of the -1 and -2 topoisomer bands. Other minor bands were not quantified.

**Supplementary Figure S7.**
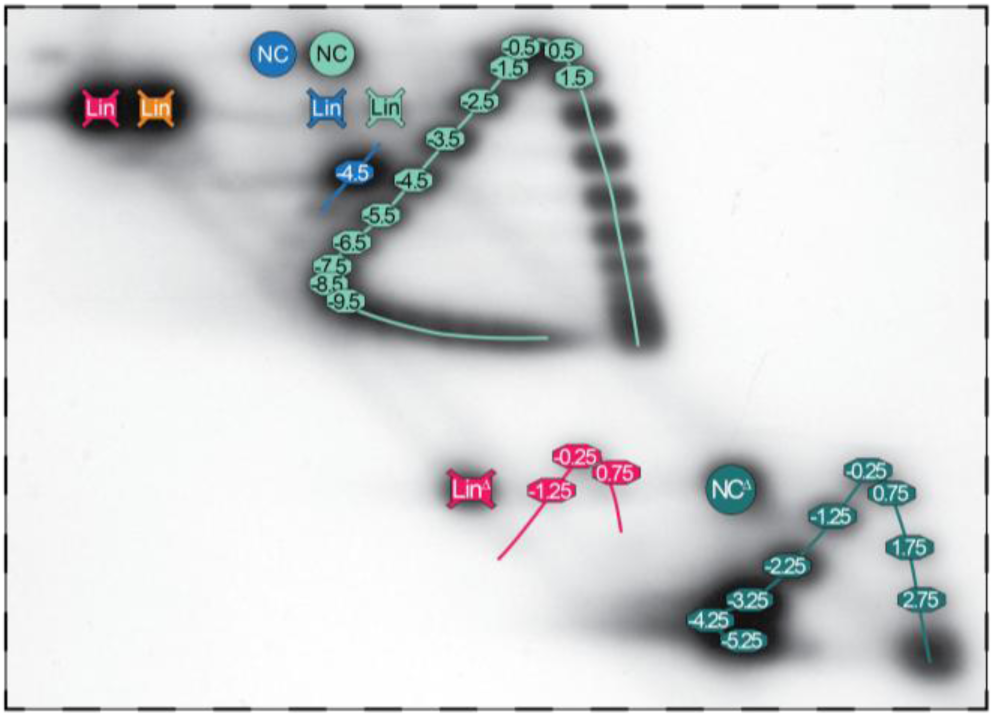
Measurement of linkage change on an evenly spaced substrate. Southern blot as shown in Figure 5e with alternative topoisomer annotations. Abbreviations: NC, nicked-circular pEVEN; NCΔ, nicked-circular pEVENΔ; Lin, linear pEVEN; LinΔ, linear pEVENΔ. Numbers correspond to topoisomer species increasing clockwise starting from the most relaxed topoisomer in the **first dimension** of electrophoresis. The isolated Xer substrate topoisomer A (blue) and its pEVENΔ recombination products (red) are numbered by comparison to the ladders (pEVEN, light green; pEVENΔ, dark green).

**Supplementary Figure S8.**
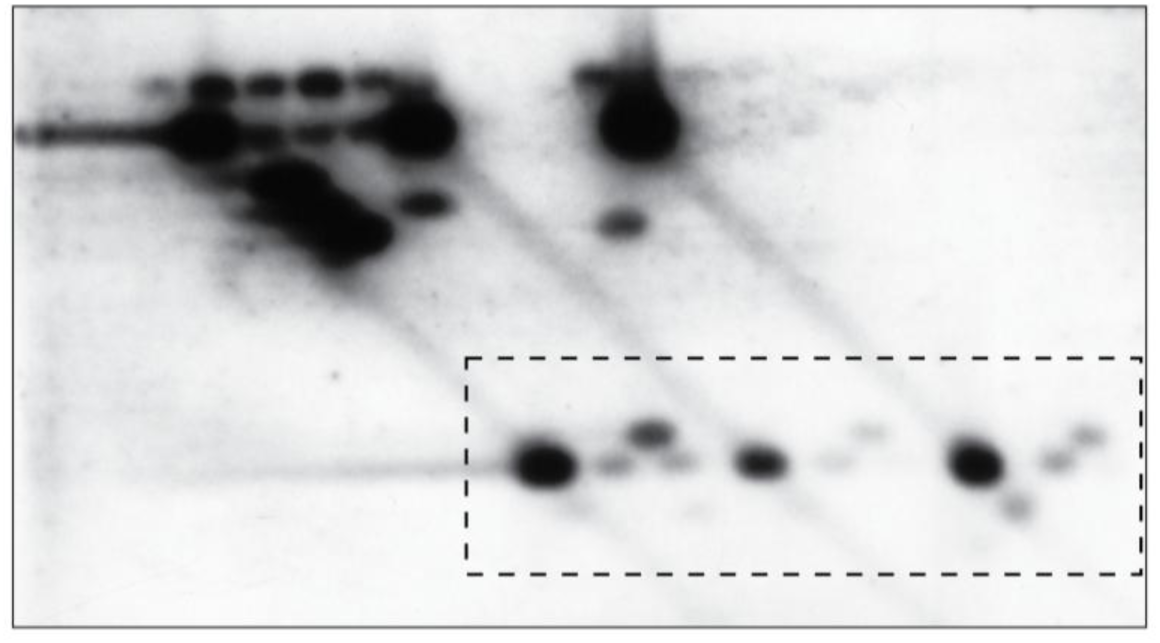
pEVENΔ product shifts. Overexposed version of the 2D southern blot shown in Figure 6b-c. Dashed lines indicate the area where this overexposed version was used as insets for Figure 6b-c to improve visibility of low concentration DNA species.

**Supplementary Figure S9.**
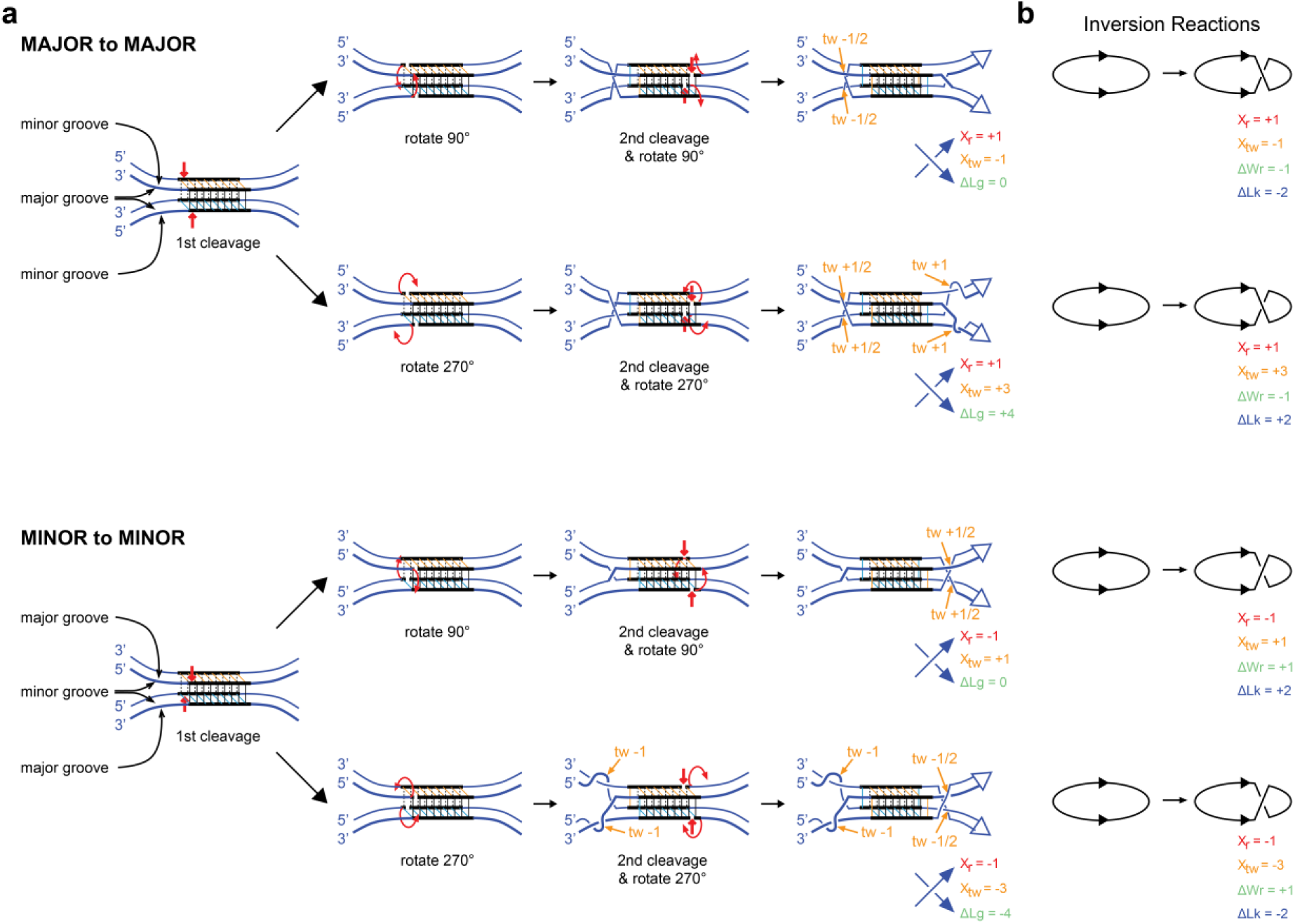
a. Four-strand parallel HJ models for strand exchange. Tyrosine recombinases cleave their recombination sites at staggered positions on either side of homologous 6-8 bp spacer regions to produce 5’OH and 3’ phosphotyrosines. Early models for recombination by these recombinases proposed synapsis by parallel alignment of recombination core sites with the four strands arranged in a square ^58,59,78^. These models proposed DNA-DNA recognition by non-Watson Crick base pairing in these 4-strand intermediates, but the topological outcome does not depend on this. Core sites can be aligned either with major-grooves (top) or minor-grooves (bottom) oriented towards each other. The leftmost diagram shows cleavage of the first pair of strands (red vertical arrows). Strand exchange then occurs by rotation of the cleaved strands around the uncleaved strands outside the spacer region by either 90° or 270° (curved red arrows). Cleavage of the second strand at the other end of the spacer (red vertical arrow) and an equivalent rotation to exchange second strands produces recombinant products. The sign of the crossing introduced between the sites (X_r_) and the total twist introduced into the sites (X_tw_) depends on whether the sites are aligned minor groove to minor groove, or major groove to major groove, and whether strand exchange occurs by rotation of 90° or 270°. All four possible outcomes are shown. The helical nature of the DNA has been removed for simplicity. **b. Predicted linkage change (DLk) for inversion reactions yielding unknotted circles.** As the sites are in inverted (head-to-head) repeat in the circular substrate, the writhe introduced by strand exchange (DWr) has the opposite sign to the local crossing introduced between the two recombination core sites (X_r_). Depending on whether the sites are juxtaposed minor-groove to minor-groove, or major-groove to major-groove and the direction of strand rotation, DLk is + or – 2 as shown, in agreement with the observed +/-2 linkage change for these reactions.

**Supplementary Figure S10.**
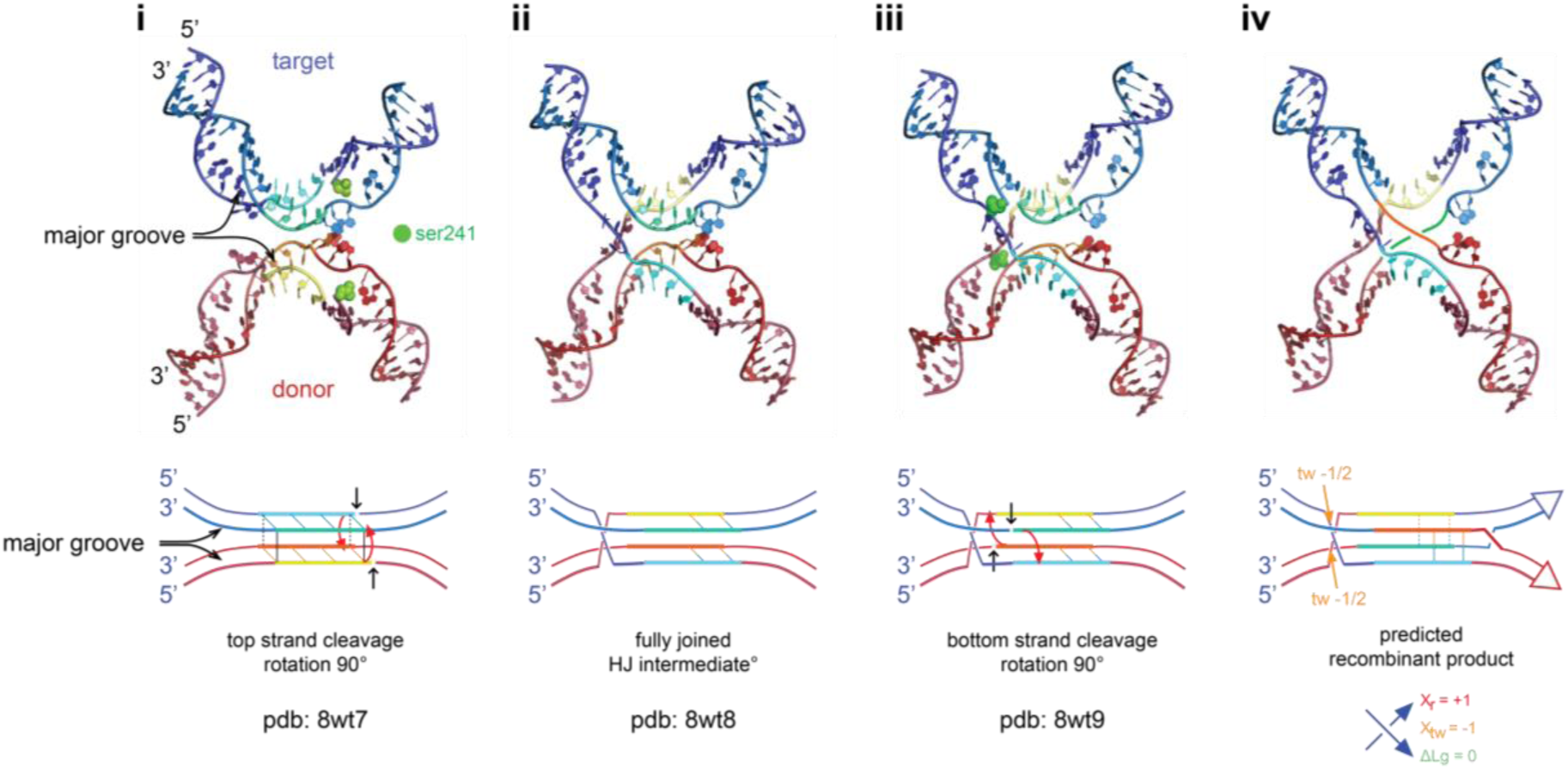
Predicted linkage change for bridge RNA-guided recombinases. Models for the bridge-RNA-guided recombinase strand exchange mechanism based on cryo electron microscopy structures of IS110 recombinase in complex with donor and target DNA at three different stages of the reaction ^27^. Bridge RNA-guided recombinases are thought to cleave DNA with a 4-nucleotide stagger to produce overhanging 3’ OH ends, and recessed 5’ ends covalently linked to the recombinase via 5’ phosphoserine linkages. **i**, Prior to strand exchange but after cleavage of top strands (pdb: 8wt7). DNA strands are base-paired to the bridge RNA and significantly underwound. Removal of the protein and bridge RNA from the structural model reveals that the two DNA sites are aligned in parallel with major grooves oriented towards each other, equivalent to the alignment shown for tyrosine recombinases in supplementary Figure S9a. Serine 241 residues from two different recombinase protomers are covalently linked to top strands at the right of each 4-nucleotide spacer. **ii**, Swapping of spacer top strands by a 90° rotation of cleaved strands around uncleaved strands followed by nucleophilic attack of the 3’ OH on the phosphoserine produces a fully ligated HJ with recombinant strands forming a right-handed (+) crossing (pdb: 8wt8). **iii**, Cleavage of the bottom strands at the other ends of the spacers by serine 241 residues from the second pair of recombinase protomers (pdb: 8wt9). **iv**, Proposed swapping of the spacer bottom strands would yield recombinant product with the two sites forming a right-handed (+) crossing. There is no structural data to support this final step. The 90° rotations introduce a half twist into each site. The overall topology of the mechanism in this projection is P = (0) parallel, R = (+1), X_r_ = +1, X_tw_ = -1 and ΔLg = 0. Top, recombination site DNA is shown in cartoon representation. Protein and bridge RNA have been removed for clarity, serine 241 is represented as green spheres. Images prepared using Pymol ^91^. Bottom, 4-strand representations of the DNA are shown, analogous to the representations in Supplementary Figure S9. Top and bottom strands of the target DNA are shown as Slate and Marine blue, top and bottom strands of the donor DNA are shown as Raspberry and Brick red. The 4-nucleotide overlap sequences are shown in cyan (target top strand), blue-green (target bottom strand), yellow (donor top strand) and orange (donor bottom strand). The same colour code is used in top and bottom panels. In **iv** the proposed path of the spacer bottom strands are drawn as lines using the same colour code. Overlap sequences are base-paired to the bridge RNA and not to each other, but base complementarity between strands in parental and recombinant configurations is indicated by horizontal sticks. Note that only the CpT / ApG dinucleotides at the left of the spacer match in recombinant products.

**Supplementary Figure S11.**
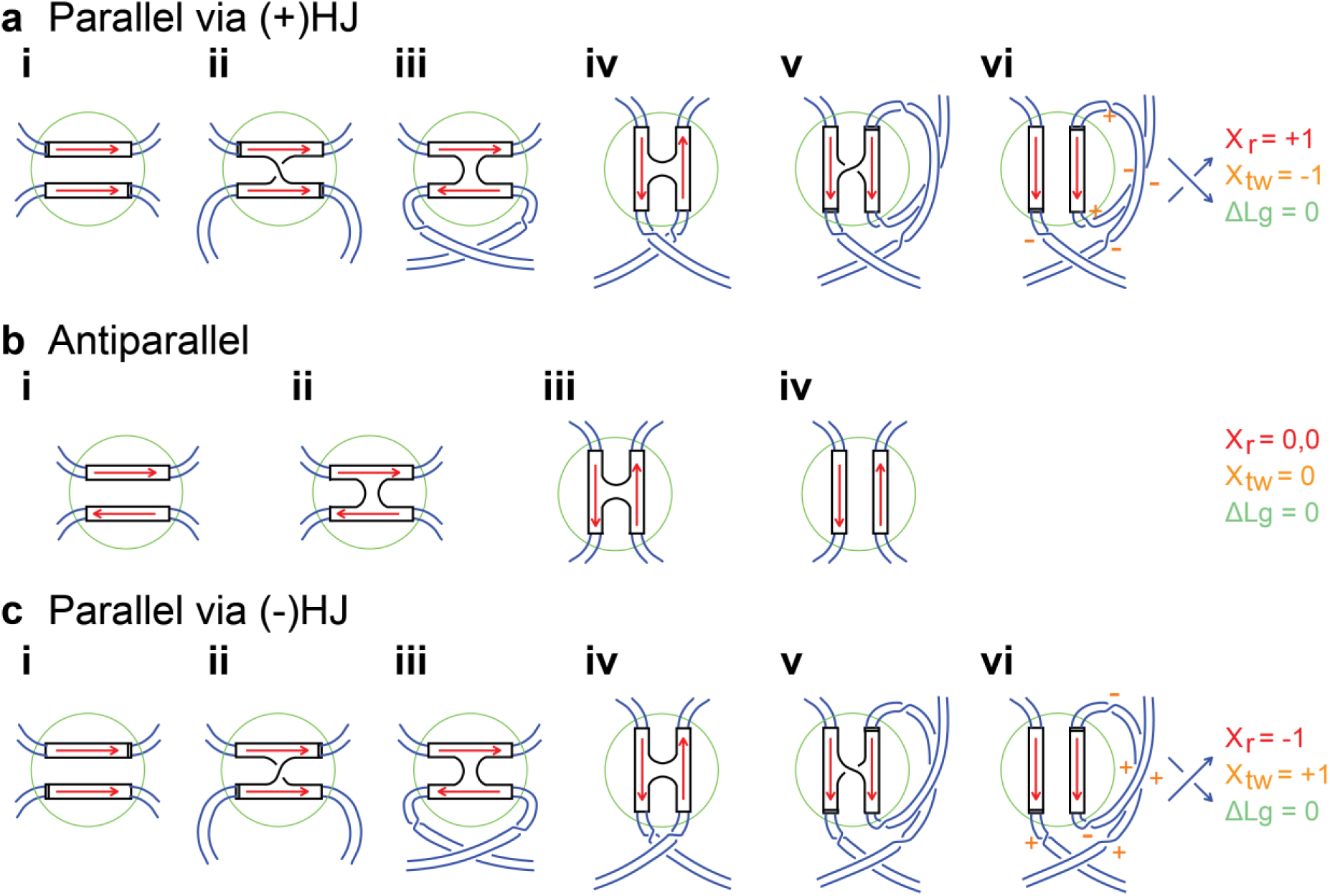
Topological outcomes of symmetrical strand exchange models via parallel or antiparallel intermediates. Red arrows indicate the direction of core sites. Regions within green circles show local movement of sites. Movement of connecting DNA strands (blue lines) outside are shown to keep track of “global” changes in twist and overall crossings between the two sites. The resultant global topological changes are summarised by values of X_r_, X_tw_ and ΔLg for each model. **a**, Parallel alignment of sites and recombination via HJs with (+) crossed strands. **i,** Sites are aligned in parallel. **ii**, first strand exchange produces a (+) HJ. **iii**, anticlockwise rotation of one site by 180° aligns the sites in antiparallel. **iv**, sites isomerise to put the other pair of strands into a crossed configuration. **v**, clockwise rotation to produce a (+) HJ. **vi**, second strand exchange to produce the recombinant product. Orange “+” and “-” signs indicate half-twists introduced by the strand exchange mechanism. **b**, Antiparallel alignment of sites and strand exchange with minimal movement of DNA ends. **i**, Sites are aligned in parallel. **ii**, strand exchange produces an antiparallel HJ that may be stacked (as shown) or square planar. **iii**, minimal isomerisation leaves the other pair of strands crossed. **iv**, exchange of the second pair of strands produces recombinant product with antiparallel sites in the R = (0,0) configuration. **c**, Parallel alignment of sites and recombination via (-) crossed HJs. **i**, Sites are aligned in parallel. **ii**, first strand exchange produces a (-) HJ. **iii**, clockwise rotation of one site by 180° aligns the sites in antiparallel. **iv**, sites isomerise to put the other pair of strands into a crossed configuration. **v**. anticlockwise rotation to produce a (-) HJ. **vi**, second strand exchange to produce the recombinant product.

**Supplementary Figure S12.**
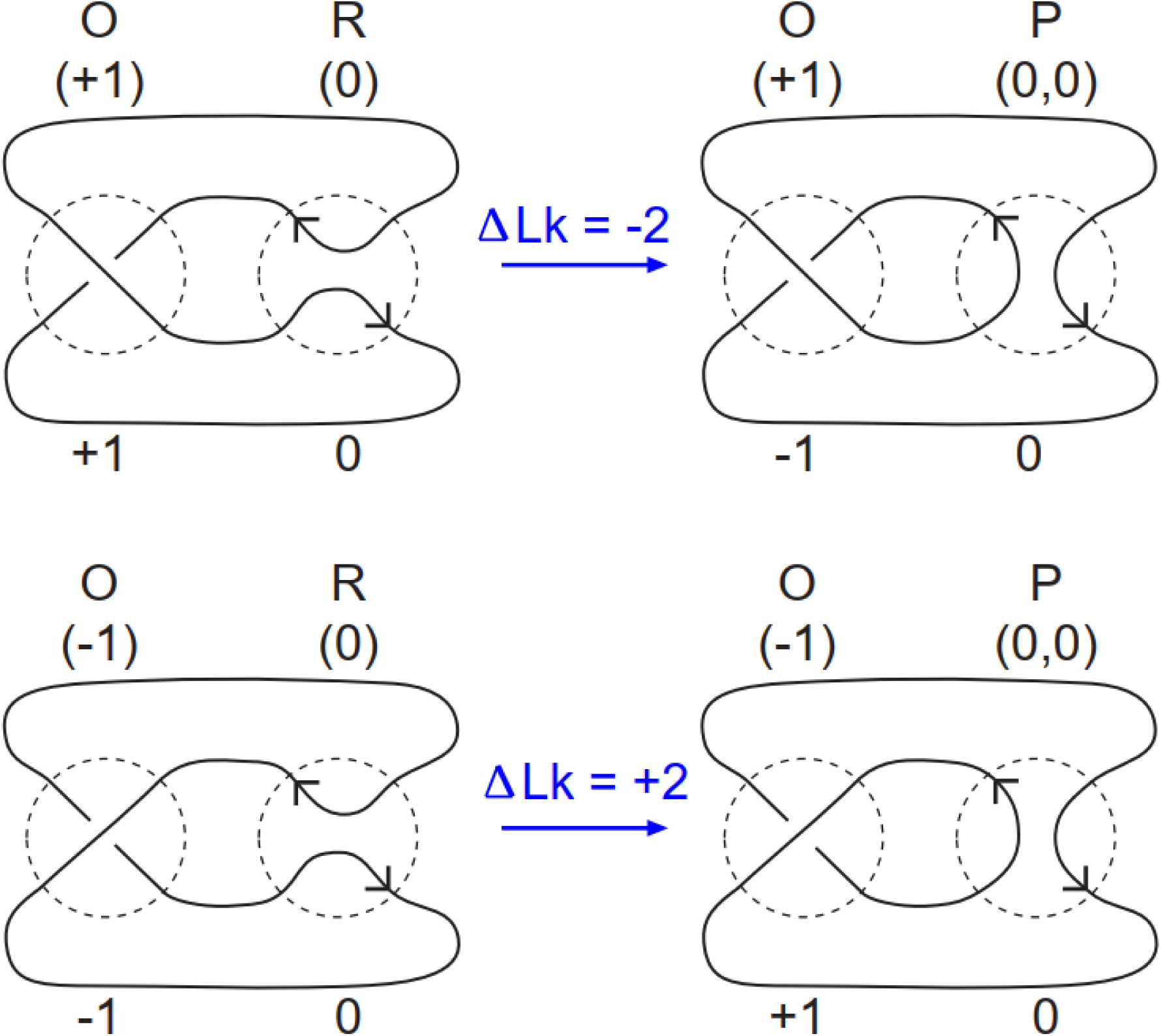
Antiparallel tangle models showing ΔLk for lambda integrase *attL* x *attR* (excisive) recombination. Tangle diagrams (within dashed circles) are shown for O (combining O_b_ and O_f_), P and R with tangle notations in brackets above each diagram. The contribution of each tangle to the linkage of each molecule is shown below each tangle. Both models show the parental antiparallel (0) tangle P being converted to the recombinant (0, 0) R tangle by the recombination mechanism. A single crossing is required in O to allow the sites to align in antiparallel. The linkage associated with O changes sign because of the change in connectivity during recombination, while the O tangle itself remains unchanged. If strand exchange occurs with no net change in linkage, as is the case if X_tw_ = 0, then ΔLk is either +2 or -2 as shown. Reactions could also occur in the reverse direction with inversion of the sign of ΔLk.

**Supplementary Table 1.**
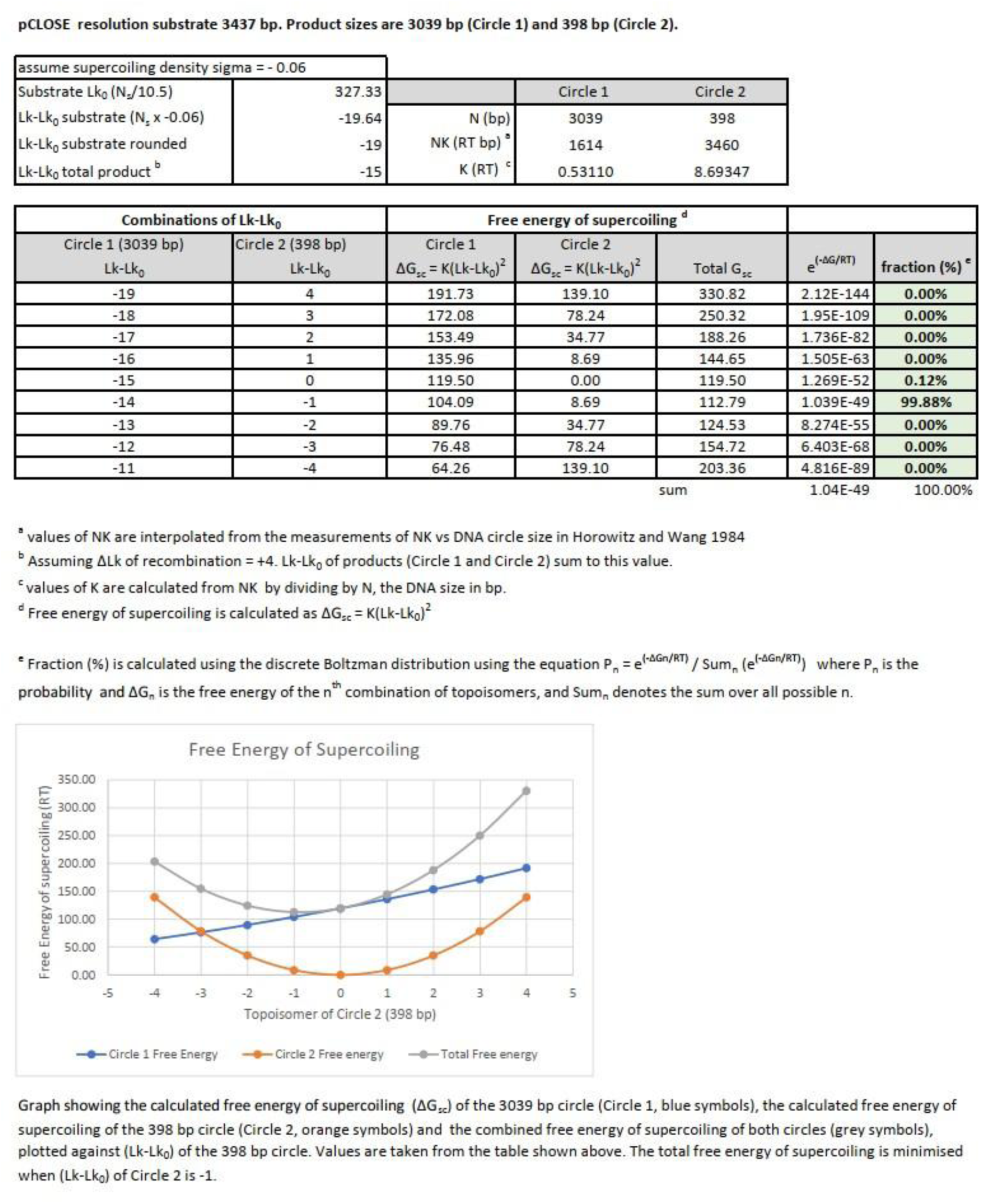
: Predicted segregation of supercoiling between the two circular products of pCLOSE

**Supplementary Table 2.**
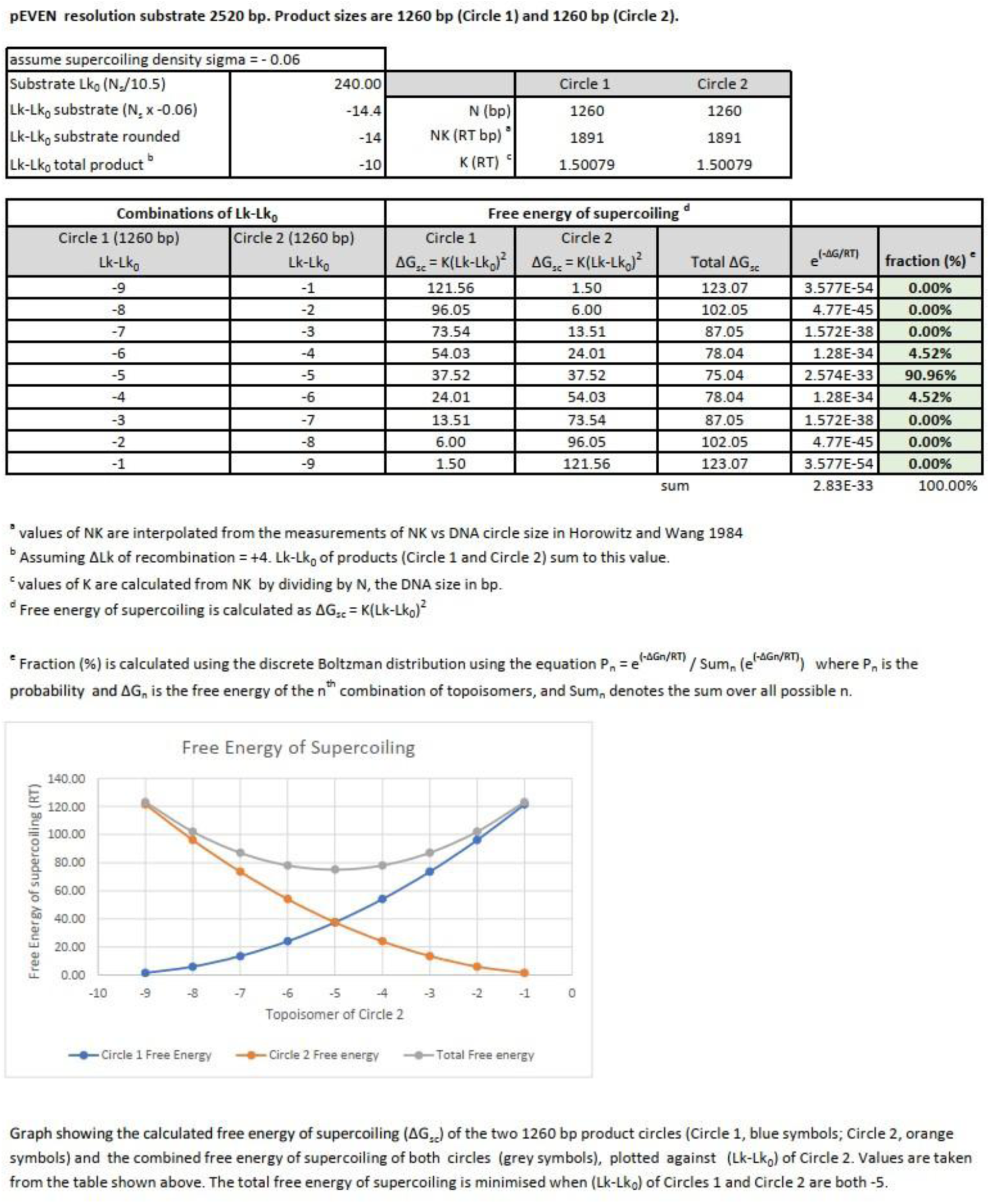
: Predicted segregation of supercoiling between the two circular products of pEVEN

## References

1. Midonet, C., and Barre, F.-X. (2014). Xer site-specific recombination: promoting vertical and horizontal transmission of genetic information. Microbiol. Spectr. 2. 10.1128/microbiolspec.mdna3-0056-2014.

2. Landy, A. (2015). The ë integrase site-specific recombination pathway. Microbiol. Spectr. 3. 10.1128/microbiolspec.MDNA3-0051-2014.

3. Blanchais, C., Pages, C., Campos, M., Boubekeur, K., Contarin, R., Orlando, M., Siguier, P., Laaberki, M.-H., Cornet, F., Charpentier, X., et al. (2025). Interplay between the Xer recombination system and the dissemination of antibioresistance in *Acinetobacter baumannii*. Nucleic Acids Res. 53, gkae1255. 10.1093/nar/gkae1255.

4. Merino, M., Acosta, J., Poza, M., Sanz, F., Beceiro, A., Chaves, F., and Bou, G. (2010). OXA-24 carbapenemase gene flanked by XerC/XerD-like recombination sites in different plasmids from different *Acinetobacter* species isolated during a nosocomial outbreak. Antimicrob. Agents Chemother. 54, 2724–2727. 10.1128/AAC.01674-09.

5. Mindlin, S., Beletsky, A., Mardanov, A., and Petrova, M. (2019). Adaptive dif modules in permafrost strains of *Acinetobacter lwoffii* and their distribution and abundance among present day *Acinetobacter* strains. Front. Microbiol. 10, 1–11. 10.3389/fmicb.2019.00632.

6. Shao, M., Ying, N., Liang, Q., Ma, N., Leptihn, S., Yu, Y., Chen, H., Liu, C., and Hua, X. (2023). P*dif*-mediated antibiotic resistance genes transfer in bacteria identified by pdifFinder. Brief. Bioinform. 24, bbac521. 10.1093/bib/bbac521.

7. Jayaram, M., Ma, C.-H., Kachroo, A.H., Rowley, P.A., Guga, P., Fan, H.-F., and Voziyanov, Y. (2015). An overview of tyrosine site-specific recombination: from an Flp perspective. Microbiol. Spectr. 3. 10.1128/microbiolspec.mdna3-0021-2014.

8. Stark, W.M. (2014). The serine recombinases. Microbiol. Spectr. 2. 10.1128/microbiolspec.mdna3-0046-2014.

9. Johnson, R.C. (2007). Bacterial site-specific DNA inversion systems. In Mobile DNA II (John Wiley & Sons, Ltd), pp. 230–271. 10.1128/9781555817954.ch13.

10. Johnson, R.C. (2015). Site-specific DNA inversion by serine recombinases. Microbiol. Spectr. 3, 1–36. 10.1128/microbiolspec.MDNA3-0047-2014.

11. Escudero, J.A., Loot, C., Nivina, A., and Mazel, D. (2015). The integron: adaptation on demand. Microbiol. Spectr. 3. 10.1128/microbiolspec.MDNA3-0019-2014.

12. Hickman, A.B., and Dyda, F. (2015). Mechanisms of DNA transposition. Microbiol. Spectr. 3. 10.1128/microbiolspec.mdna3-0034-2014.

13. Castillo, F., Benmohamed, A., and Szatmari, G. (2017). Xer site specific recombination: double and single recombinase systems. Front. Microbiol. 8, 1–18. 10.3389/fmicb.2017.00453.

14. Colloms, S.D. (2013). The topology of plasmid-monomerizing Xer site-specific recombination. Biochem. Soc. Trans. 41, 589–594. 10.1042/BST20120340.

15. Van Duyne, G.D. (2015). Cre recombinase. Microbiol. Spectr. 3. 10.1128/microbiolspec.mdna3-0014-2014.

16. Meinke, G., Bohm, A., Hauber, J., Pisabarro, M.T., and Buchholz, F. (2016). Cre recombinase and other tyrosine recombinases. Chem. Rev. 116, 12785–12820. 10.1021/acs.chemrev.6b00077.

17. Grainge, I. (2013). Simple topology: FtsK-directed recombination at the *dif* site. Biochem. Soc. Trans. 41, 595–600. 10.1042/BST20120299.

18. Cadden, G.M., Schloetel, J.-G., McKenzie, G., Boocock, M.R., Magennis, S.W., and Stark, W.M. (2024). Direct observation of subunit rotation during DNA strand exchange by serine recombinases. Nat. Commun. 15, 10407. 10.1038/s41467-024-54531-4.

19. Stark, W.M., Sherratt, D.J., and Boocock, M.R. (1989). Site-specific recombination by Tn3 resolvase: topological changes in the forward and reverse reactions. Cell 58, 779–790. 10.1016/0092-8674(89)90111-6.

20. Olorunniji, F.J., Buck, D.E., Colloms, S.D., McEwan, A.R., Smith, M.C.M., Stark, W.M., and Rosser, S.J. (2012). Gated rotation mechanism of site-specific recombination by ϕC31 integrase. Proc. Natl. Acad. Sci. U. S. A. 109, 19661–19666. 10.1073/pnas.1210964109.

21. Li, W., Kamtekar, S., Xiong, Y., Sarkis, G.J., Grindley, N.D.F., and Steitz, T.A. (2005). Structure of a synaptic ãä resolvase tetramer covalently linked to two cleaved DNAs. Science 309, 1210–1215. 10.1126/science.1112064.

22. Stachowski, K., Norris, A.S., Potter, D., Wysocki, V.H., and Foster, M.P. (2022). Mechanisms of Cre recombinase synaptic complex assembly and activation illuminated by Cryo-EM. Nucleic Acids Res. 50, 1753–1769. 10.1093/nar/gkac032.

23. Arciszewska, L.K., Grainge, I., and Sherratt, D.J. (1997). Action of site-specific recombinases XerC and XerD on tethered Holliday junctions. EMBO J. 16, 3731–3743. 10.1093/emboj/16.12.3731.

24. Guo, F., Gopaul, D.N., and Van Duyne, G.D. (1997). Structure of Cre recombinase complexed with DNA in a site-specific recombination synapse. Nature 389, 40–46. 10.1038/37925.

25. Gopaul, D.N., Guo, F., and Van Duyne, G.D. (1998). Structure of the Holliday junction intermediate in Cre–*loxP* site-specific recombination. EMBO J. 17, 4175–4187. 10.1093/emboj/17.14.4175.

26. Azaro, M.A., and Landy, A. (1997). The isomeric preference of Holliday junctions influences resolution bias by Lambda integrase. EMBO J. 16, 3744–3755. 10.1093/emboj/16.12.3744.

27. Hiraizumi, M., Perry, N.T., Durrant, M.G., Soma, T., Nagahata, N., Okazaki, S., Athukoralage, J.S., Isayama, Y., Pai, J.J., Pawluk, A., et al. (2024). Structural mechanism of bridge RNA-guided recombination. Nature 630, 994–1002. 10.1038/s41586-024-07570-2.

28. McClintock, B. (1932). A correlation of ring-shaped chromosomes with variegation in zea mays. Proc. Natl. Acad. Sci. 18, 677–681. 10.1073/pnas.18.12.677.

29. Kuempel, P.L., Henson, J.M., Dircks, L., Tecklenburg, M., and Lim, D.F. (1991). *dif*, a recA-independent recombination site in the terminus region of the chromosome of *Escherichia coli*. New Biol. 3, 799–811.

30. Blakely, G., May, G., McCulloch, R., Arciszewska, L.K., Burke, M., Lovett, S.T., and Sherratt, D.J. (1993). Two related recombinases are required for site-specific recombination at *dif* and *cer* in *E. coli* K12. Cell 75, 351–361. 10.1016/0092-8674(93)80076-Q.

31. Colloms, S.D., McCulloch, R., Grant, K., Neilson, L., and Sherratt, D.J. (1996). Xer-mediated site-specific recombination *in vitro*. EMBO J. 15, 1172–1181. 10.1002/j.1460-2075.1996.tb00456.x.

32. McCulloch, R., Coggins, L.W., Colloms, S.D., and Sherratt, D.J. (1994). Xer-mediated site-specific recombination at *cer* generates Holliday junctions *in vivo*. EMBO J. 13, 1844–1855. 10.1002/j.1460-2075.1994.tb06453.x.

33. Arciszewska, L.K., and Sherratt, D.J. (1995). Xer site-specific recombination *in vitro*. EMBO J. 14, 2112–2120. 10.1002/j.1460-2075.1995.tb07203.x.

34. Colloms, S.D., Sykora, P., Szatmari, G., and Sherratt, D.J. (1990). Recombination at ColE1 *cer* requires the *Escherichia coli xerC* gene product, a member of the Lambda integrase family of site-specific recombinases. J. Bacteriol. 172, 6973–6980. 10.1128/jb.172.12.6973-6980.1990.

35. Aussel, L., Barre, F.X., Aroyo, M., Stasiak, A., Stasiak, A.Z., and Sherratt, D. (2002). FtsK is a DNA motor protein that activates chromosome dimer resolution by switching the catalytic state of the XerC and XerD recombinases. Cell 108, 195–205. 10.1016/s0092-8674(02)00624-4.

36. Barre, F.X., Aroyo, M., Colloms, S.D., Helfrich, A., Cornet, F., and Sherratt, D.J. (2000). FtsK functions in the processing of a Holliday junction intermediate during bacterial chromosome segregation. Genes Dev. 14, 2976–2988. 10.1101/gad.188700.

37. Begg, K.J., Dewar, S.J., and Donachie, W.D. (1995). A new *Escherichia coli* cell division gene, *ftsK*. J. Bacteriol. 177, 6211–6222. 10.1128/jb.177.21.6211-6222.1995.

38. Bigot, S., Saleh, O.A., Lesterlin, C., Pages, C., Karoui, M.E., Dennis, C., Grigoriev, M., Allemand, J.-F., Barre, F.-X., and Cornet, F. (2005). KOPS: DNA motifs that control *E. coli* chromosome segregation by orienting the FtsK translocase. EMBO J. 24, 3770. 10.1038/sj.emboj.7600835.

39. Bigot, S., Saleh, O.A., Cornet, F., Allemand, J.-F., and Barre, F.-X. (2006). Oriented loading of FtsK on KOPS. Nat. Struct. Mol. Biol. 13, 1026–1028. 10.1038/nsmb1159.

40. Grainge, I., Lesterlin, C., and Sherratt, D.J. (2011). Activation of XerCD-*dif* recombination by the FtsK DNA translocase. Nucleic Acids Res. 39, 5140–5148. 10.1093/nar/gkr078.

41. Summers, D.K., Beton, C.W.H., and Withers, H.L. (1993). Multicopy plasmid instability: the dimer catastrophe hypothesis. Mol. Microbiol. 8, 1031–1038. 10.1111/j.1365-2958.1993.tb01648.x.

42. Field, C.M., and Summers, D.K. (2011). Multicopy plasmid stability: revisiting the dimer catastrophe. J. Theor. Biol. 291, 119–127. 10.1016/j.jtbi.2011.09.006.

43. Summers, D.K., and Sherratt, D.J. (1984). Multimerization of high copy number plasmids causes instability: CoIE1 encodes a determinant essential for plasmid monomerization and stability. Cell 36, 1097–1103. 10.1016/0092-8674(84)90060-6.

44. Friehs, K. (2004). Plasmid copy number and plasmid stability. In New trends and developments in biochemical engineering, T. Scheper, ed. (Springer), pp. 47–82. 10.1007/b12440.

45. Cornet, F., Mortier, I., Patte, J., and Louarn, J.M. (1994). Plasmid pSC101 harbors a recombination site, *psi*, which is able to resolve plasmid multimers and to substitute for the analogous chromosomal *Escherichia coli* site *dif*. J. Bacteriol. 176, 3188–3195. 10.1128/jb.176.11.3188-3195.1994.

46. Summers, D.K., and Sherratt, D.J. (1988). Resolution of ColE1 dimers requires a DNA sequence implicated in the three-dimensional organization of the cer site. EMBO J. 7, 851–858.

47. Colloms, S.D., Bath, J., and Sherratt, D.J. (1997). Topological selectivity in Xer site-specific recombination. Cell 88, 855–864. 10.1016/S0092-8674(00)81931-5.

48. Pham, H., Dery, K.J., Sherratt, D.J., and Tolmasky, M.E. (2002). Osmoregulation of dimer resolution at the plasmid pJHCMW1 *mwr* locus by *Escherichia coli* XerCD recombination. J. Bacteriol. 184, 1607–1616. 10.1128/JB.184.6.1607-1616.2002.

49. Fournes, F., Campos, M., Cury, J., Schiavon, C., Pagès, C., Touchon, M., Rocha, E.P.C., Rousseau, P., and Cornet, F. (2025). The pathway to resolve dimeric forms distinguishes plasmids from megaplasmids in *Enterobacteriaceae*. Nucleic Acids Res. 53, gkae1300. 10.1093/nar/gkae1300.

50. Sträter, N., Sherratt, D.J., and Colloms, S.D. (1999). X-ray structure of aminopeptidase A from *Escherichia coli* and a model for the nucleoprotein complex in Xer site-specific recombination. EMBO J. 18, 4513–4522. 10.1093/emboj/18.16.4513.

51. Alén, C., Sherratt, D.J., and Colloms, S.D. (1997). Direct interaction of aminopeptidase A with recombination site DNA in Xer site-specific recombination. EMBO J. 16, 5188–5197. 10.1093/emboj/16.17.5188.

52. McCulloch, R., Burke, M.E., and Sherratt, D.J. (1994). Peptidase activity of *Escherichia coli* aminopeptidase A is not required for its role in Xer site-specific recombination. Mol. Microbiol. 12, 241–251. 10.1111/j.1365-2958.1994.tb01013.x.

53. Stirling, C.J., Colloms, S.D., Collins, J.F., Szatmari, G., and Sherratt, D.J. (1989). *xerB*, an escherichia coli gene required for plasmid ColE1 site-specific recombination, is identical to *pepA*, encoding aminopeptidase A, a protein with substantial similarity to bovine lens leucine aminopeptidase. EMBO J. 8, 1623–1627. 10.1002/j.1460-2075.1989.tb03547.x.

54. Stirling, C.J., Szatmari, G., Stewart, G., Smith, M.C., and Sherratt, D.J. (1988). The arginine repressor is essential for plasmid-stabilizing site-specific recombination at the ColE1 *cer* locus. EMBO J. 7, 4389–4395. 10.1002/j.1460-2075.1988.tb03338.x.

55. Colloms, S.D., Alén, C., and Sherratt, D.J. (1998). The ArcA/ArcB two-component regulatory system of *Escherichia coli* is essential for Xer site-specific recombination at *psi*. Mol. Microbiol. 28, 521–530. 10.1046/j.1365-2958.1998.00812.x.

56. Bregu, M., Sherratt, D.J., and Colloms, S.D. (2002). Accessory factors determine the order of strand exchange in Xer recombination at *psi*. EMBO J. 21, 3888–3897. 10.1093/emboj/cdf379.

57. Holmes, E.P., Gamill, M.C., Provan, J.I., Wiggins, L., Rusková, R., Whittle, S., Catley, T.E., Main, K.H.S., Shephard, N., Bryant, H.E., et al. (2025). Quantifying complexity in DNA structures with high resolution Atomic Force Microscopy. Nat. Commun. 16, 5482. 10.1038/s41467-025-60559-x.

58. Nash, H.A., and Pollock, T.J. (1983). Site-specific recombination of bacteriophage Lambda: the change in topological linking number associated with exchange of DNA strands. J. Mol. Biol. 170, 19–38. 10.1016/S0022-2836(83)80225-3.

59. Kikuchi, Y., and Nash, H.A. (1979). Nicking-closing activity associated with bacteriophage Lambda int gene product. Proc. Natl. Acad. Sci. 76, 3760–3764. 10.1073/pnas.76.8.3760.

60. Huffman, K.E., and Levene, S.D. (1999). DNA-sequence asymmetry directs the alignment of recombination sites in the FLP synaptic complex. J. Mol. Biol. 286, 1–13. 10.1006/jmbi.1998.2468.

61. Laxmikanthan, G., Xu, C., Brilot, A.F., Warren, D., Steele, L., Seah, N., Tong, W., Grigorieff, N., Landy, A., and Van Duyne, G.D. (2016). Structure of a Holliday junction complex reveals mechanisms governing a highly regulated DNA transaction. eLife 5, e14313. 10.7554/eLife.14313.

62. Bebel, A., Karaca, E., Kumar, B., Stark, W.M., and Barabas, O. (2016). Structural snapshots of Xer recombination reveal activation by synaptic complex remodeling and DNA bending. eLife 5, e19706. 10.7554/eLife.19706.

63. Shoura, M.J., Giovan, S.M., Vetcher, A.A., Ziraldo, R., Hanke, A., and Levene, S.D. (2020). Loop-closure kinetics reveal a stable, right-handed DNA intermediate in Cre recombination. Nucleic Acids Res. 48, 4371–4381. 10.1093/nar/gkaa153.

64. Crisona, N.J., Weinberg, R.L., Peter, B.J., Sumners, D.W., and Cozzarelli, N.R. (1999). The topological mechanism of phage Lambda integrase. J. Mol. Biol. 289, 747–775. 10.1006/jmbi.1999.2771.

65. Stark, W.M., and Boocock, M.R. (1994). The linkage change of a knotting reaction catalysed by Tn3 resolvase. J. Mol. Biol. 239, 25–36. 10.1006/jmbi.1994.1348.

66. Lutz, C.T., Hollifield, W.C., Seed, B., Davie, J.M., and Huang, H.V. (1987). Syrinx 2A: an improved Lambda phage vector designed for screening DNA libraries by recombination *in vivo*. Proc. Natl. Acad. Sci. U. S. A. 84, 4379–4383. 10.1073/pnas.84.13.4379.

67. Seed, B. (1983). Purification of genomic sequences from bacteriophage libraries by recombination and selection *in vivo*. Nucleic Acids Res. 11, 2427–2445. 10.1093/nar/11.8.2427.

68. Summers, D.K., and Sherratt, D.J. (1988). Resolution of ColE1 dimers requires a DNA sequence implicated in the three-dimensional organization of the *cer* site. EMBO J. 7, 851–858. 10.1002/j.1460-2075.1988.tb02884.x.

69. Subramanya, H.S., Arciszewska, L.K., Baker, R.A., Bird, L.E., Sherratt, D.J., and Wigley, D.B. (1997). Crystal structure of the site-specific recombinase, XerD. EMBO J. 16, 5178–5187. 10.1093/emboj/16.17.5178.

70. Sambrook, J., and Russell, D.W. (2001). Molecular cloning: a laboratory manual (CSHL Press).

71. Schindelin, J., Arganda-Carreras, I., Frise, E., Kaynig, V., Longair, M., Pietzsch, T., Preibisch, S., Rueden, C., Saalfeld, S., Schmid, B., et al. (2012). Fiji: an open-source platform for biological-image analysis. Nat. Methods 9, 676–682. 10.1038/nmeth.2019.

72. Balagurumoorthy, P., Adelstein, S.J., and Kassis, A.I. (2008). Method to eliminate linear DNA from mixture containing nicked circular, supercoiled, and linear plasmid DNA. Anal. Biochem. 381, 172–174. 10.1016/j.ab.2008.06.037.

73. Irobalieva, R.N., Fogg, J.M., Catanese, D.J., Sutthibutpong, T., Chen, M., Barker, A.K., Ludtke, S.J., Harris, S.A., Schmid, M.F., Chiu, W., et al. (2015). Structural diversity of supercoiled DNA. Nat. Commun. 6, 1–10. 10.1038/ncomms9440.

74. Fogg, J.M., Kolmakova, N., Rees, I., Magonov, S., Hansma, H., Perona, J.J., and Zechiedrich, E.L. (2006). Exploring writhe in supercoiled minicircle DNA. J. Phys. Condens. Matter Inst. Phys. J. 18, S145–S159. 10.1088/0953-8984/18/14/S01.

75. Horowitz, D.S., and Wang, J.C. (1984). Torsional rigidity of DNA and length dependence of the free energy of DNA supercoiling. J. Mol. Biol. 173, 75–91. 10.1016/0022-2836(84)90404-2.

76. Shore, D., and Baldwin, R.L. (1983). Energetics of DNA twisting. II. Topoisomer analysis. J. Mol. Biol. 170, 983–1007. 10.1016/s0022-2836(83)80199-5.

77. Wang, J.C. (1980). Superhelical DNA. Trends Biochem. Sci. 5, 219–221. 10.1016/S0968-0004(80)80012-0.

78. Abremski, K., Frommer, B., and Hoess, R.H. (1986). Linking-number changes in the DNA substrate during Cre-mediated *loxP* site-specific recombination. J. Mol. Biol. 192, 17–26. 10.1016/0022-2836(86)90460-2.

79. Grainge, I., Buck, D., and Jayaram, M. (2000). Geometry of site alignment during Int family recombination: Antiparallel synapsis by the Flp recombinase. J. Mol. Biol. 298, 749–764. 10.1006/jmbi.2000.3679.

80. Sumners, D.W., Ernst, C., Spengler, S.J., and Cozzarelli, N.R. (1995). Analysis of the mechanism of DNA recombination using tangles. Q. Rev. Biophys. 28, 253–313. 10.1017/s0033583500003498.

81. Biswas, T., Aihara, H., Radman-Livaja, M., Filman, D., Landy, A., and Ellenberger, T. (2005). A structural basis for allosteric control of DNA recombination by λ integrase. Nature 435, 1059–1066. 10.1038/nature03657.

82. Chen, Y., Narendra, U., Iype, L.E., Cox, M.M., and Rice, P.A. (2000). Crystal structure of a Flp recombinase–Holliday junction complex: assembly of an active oligomer by helix swapping. Mol. Cell 6, 885–897. 10.1016/S1097-2765(05)00088-2.

83. Bath, J. (1997). Xer-mediated site-specific recombination *in vitro*. Thesis.

84. Vazquez, M., Colloms, S.D., and Sumners, D.W. (2005). Tangle analysis of Xer recombination reveals only three solutions, all consistent with a single three-dimensional topological pathway. J. Mol. Biol. 346, 493–504. 10.1016/j.jmb.2004.11.055.

85. Cozzarelli, N.R., Krasnow, M.A., Gerrard, S.P., and White, J.H. (1984). A topological treatment of recombination and topoisomerases. Cold Spring Harb. Symp. Quant. Biol. 49, 383–400. 10.1101/SQB.1984.049.01.045.

86. Griffith, J.D., and Nash, H.A. (1985). Genetic rearrangement of DNA induces knots with a unique topology: implications for the mechanism of synapsis and crossing-over. Proc. Natl. Acad. Sci. U. S. A. 82, 3124–3128. 10.1073/pnas.82.10.3124.

87. McGavin, S. (1989). Four strand recombination models. J. Theor. Biol. 136, 135–150. 10.1016/S0022-5193(89)80221-8.

88. Nunes-Düby, S.E., Azaro, M.A., and Landy, A. (1995). Swapping DNA strands and sensing homology without branch migration in λ site-specific recombination. Curr. Biol. 5, 139–148. 10.1016/S0960-9822(95)00035-2.

89. Mouw, K.W., Rowland, S.-J., Gajjar, M.M., Boocock, M.R., Stark, W.M., and Rice, P.A. (2008). Architecture of a serine recombinase-DNA regulatory complex. Mol. Cell 30, 145–155. 10.1016/j.molcel.2008.02.023.

90. Vanhooff, V., Galloy, C., Agaisse, H., Lereclus, D., Révet, B., and Hallet, B. (2006). Self-control in DNA site-specific recombination mediated by the tyrosine recombinase TnpI. Mol. Microbiol. 60, 617–629. 10.1111/j.1365-2958.2006.05127.x.

91. Schrödinger, Llc (2015). The PyMOL molecular graphics system, version 1.8.

